# Microbial community diversity predicts invasion resistance of freshwater biofilms against antibiotic-resistant bacteria

**DOI:** 10.64898/2026.08.04.742497

**Authors:** Elisa C. P. Catão, Uli Klümper, Giulia Gionchetta, Xavier Bellanger, Amélie Porteu de la Morandière, Kenyum Bagra, Irina Dielacher, Alan Xavier Elena, Eda Deniz Erdem, Sonia Galazka, Agata Goryluk-Salmonowicz, Mateusz Szadziul, Edina Szekeres, Adela Teban-Man, Cristian Coman, Norbert Kreuzinger, Magdalena Popowska, Julia Vierheilig, Shauna O’Shea Boland, Fiona Walsh, Markus Woegerbauer, Helmut Bürgmann, Thomas U. Berendonk, Christophe Merlin

**Affiliations:** Université de Lorraine, CNRS, LCPME, UMR 7564, Nancy, France; Université de Toulon, MAPIEM, Toulon, France; Technische Universität Dresden, Institute for Hydrobiology, Zellescher Weg 40, 01217 Dresden, Germany; Eawag, Swiss Federal Institute of Aquatic Science and Technology, Department of Surface Waters – Research and Management, 6047 Kastanienbaum, Switzerland; Institute of Environmental Assessment and Water Research (IDAEA), Spanish Council of Scientific Research (CSIC), 08034 Barcelona, Spain; Department of Civil Engineering, IIT-BHU, Banaras Hindu University Campus, Near Lanka, 221005 Varanasi, Uttar Pradesh, India; Institute of Water Quality and Resource Management, TU Wien, Karlsplatz 13/2261, 1040 Vienna, Austria; Department for Integrative Risk Assessment, Division for Risk Assessment, Data and Statistics, AGES – Austrian Agency for Health and Food Safety, Spargelfeldstraße 191, 1220 Vienna, Austria; University of Warsaw, Faculty of Biology, Institute of Microbiology, Department of Bacterial Physiology, Miecznikowa 1, 02-096 Warsaw, Poland; Department of Biology, Kathleen Lonsdale Institute for Human Health, Maynooth University, Maynooth, Co. Kildare, Ireland; Institute of Biological Research Cluj, National Institute of Research and Development for Biological Sciences, 48 Republicii Street, Cluj-Napoca 400015, Romania; Interuniversity Cooperation Centre Water & Health, Austria; Warsaw University of Life Sciences, Institute of Biology, Department of Biochemistry and Microbiology, Warsaw, Poland

**Author notes:** Contributed equally to this work (Elisa C. P. Catão, Uli Klümper, Giulia Gionchetta). Corresponding author <u>Corresponding author address:</u>.

**Keywords:** Biotic resistance, Ecological prediction, River biofilms, Phylogenetic niche occupancy, Antibiotic-resistant bacteria

## Abstract

Rivers receive continuous inputs of antibiotic-resistant bacteria (ARB) from wastewater, agriculture, and other anthropogenic sources, yet it remains unclear whether the recipient ecological component and its microbial communities determine whether introduced ARB establish or disappear. Ecological invasion theory predicts that invasion success depends on biodiversity, community stability, and occupation of ecological niche space, but these mechanisms have rarely been evaluated together in natural microbial communities. Here, we challenged river biofilms collected from 20 sites in 12 European rivers across six countries with a model antibiotic-resistant *Escherichia coli* carrying a conjugative IncP-1α plasmid The invasion assays were carried out under standardized laboratory flume conditions. River biofilms differed markedly in their permissiveness to invasion despite identical invasion conditions. Higher bacterial diversity consistently accelerated invader loss rates, whereas communities containing more abundant and diverse close phylogenetic neighbours of the invader exhibited stronger exclusion during early biofilm establishment. Diversity loss during transition into the experimental system emerged as the strongest explanatory variable of invasion resistance prior to biofilm maturation, whereas Shannon diversity became the dominant predictor in mature communities. Integrating these complementary ecological dimensions substantially improved explanatory prediction of ARB persistence compared with individual predictors alone. Particularly invasion-resistant biofilms also exhibited distinct ecological community composition consistent with mature, structurally complex microbial assemblages. Together, our findings demonstrate that the establishment of ARB in the environment is not stochastic but can be predicted from measurable ecological properties of recipient microbiomes, highlighting microbial biodiversity and community organization as natural barriers to antimicrobial resistance dissemination.

## Introduction

Antibiotic resistance is an increasing societal health challenge across human, animal, and environmental spheres [1]. Its spread is driven not only by selection during antimicrobial use [2], but also by dissemination of antibiotic-resistant bacteria (ARB) and antibiotic resistance genes (ARGs) across One Health compartment boundaries [3, 4]. Aquatic ecosystems are central to this process because water connects human activities and receives continuous inputs from wastewater, agriculture, and urban runoff [5–7].

Most efforts to understand environmental antimicrobial resistance (AMR) dissemination have focused on release pathways, selective pollutants, and genetic mechanisms promoting ARG persistence, including wastewater discharge, manure application, horizontal gene transfer, and (co-)selective compounds [8–13]. In contrast, less attention has been given to the receiving environment as an active ecological filter that determines the residence time of incoming ARB. This is important because bacteria from human and animal sources are often expected to persist poorly in natural environments [14]. Resident communities may therefore constrain AMR dissemination by rapidly excluding invading ARB or, conversely, facilitate it by allowing sufficient persistence for ARB adaptation and growth, or by horizontal transfer of ARG-carrying mobile genetic elements into the resident microbiome.

Resident microbiomes have indeed been shown to act as ecological barriers to AMR establishment under some circumstances, rather than serving only as passive recipients of ARB and ARGs. Across low-impact European environments, higher microbial diversity was associated with reduced ARG accumulation in structured soils, although comparable long-term relationships were not evident in more dynamic riverbed microbiomes [15]. Mechanistic experiments support this diversity-barrier concept: dilution-to-extinction of soil microbial communities increased the spread of AMR [16], and metal-induced biodiversity loss in river biofilms increased persistence of a model resistant *Escherichia coli* invader [17]. Similarly, wastewater stabilisation systems showed that endogenous microbiome dynamics can limit the establishment of entering ARB and ARGs [18]. Together, these findings suggest that ARB persistence in environmental communities can be understood as a biological invasion process shaped by resident microbiome structure, diversity, and stability [19, 20].

Microbial invasion is commonly described as a sequence of introduction, establishment, growth or spread, and impact on the resident community [19]. While introduction is strongly influenced by propagule pressure and invader traits [20, 21], establishment directly depends on the permissiveness or biotic resistance of the receiving community [22]. Biotic resistance theory predicts that more diverse communities should be less likely invaded because they occupy a greater range of ecological niches, deplete resources more efficiently, and increase the likelihood of competitive or antagonistic interactions [16, 19, 20, 23, 24]. Beyond overall diversity, invasion resistance may depend on whether resident taxa already occupy ecological space similar to the invader. Under phylogenetic limiting similarity and related niche-overlap concepts, close relatives or functionally similar taxa can reduce establishment by competing for similar resources or occupying comparable microhabitats [22, 25, 26]. Community disturbance and restructuring may further favor permissiveness to invasion, because loss of resident taxa or destabilisation of interaction networks can open ecological niche space and weaken priority effects or competitive exclusion [27–30]. However, while diversity, phylogenetic relatedness, and disturbance-associated restructuring have each been tested as individual determinants of microbial invasion resistance, they have rarely been integrated to quantify their relative contributions or to assess whether ARB invasion success in complex environmental microbiomes is predictable and explainable. Such predictability is essential for moving environmental AMR risk assessment beyond exposure-based measures alone.

Here, we tested whether river biofilm communities differ in their resistance to invasion by a model ARB, and whether this resistance can be predicted from measurable ecological community properties. River biofilms were chosen because they represent resident, spatially structured components of freshwater ecosystems, in contrast to more transient planktonic fractions [31]. Natural biofilms were grown on artificial supports at 20 sites across 12 rivers in 6 European countries, transferred into well-established standardised laboratory recirculating flume systems [17, 32–34] and challenged with model invader *E. coli* strain carrying the conjugative IncP-1 antibiotic resistance plasmid pG527. We monitored biofilm community dynamics using 16S rRNA gene sequencing and invader strain and plasmid persistence using qPCR. We hypothesised that ARB persistence in freshwater biofilms is not stochastic, but predictable and explainable from three ecological layers: resident biodiversity, community stability, and occupation of the invader’s phylogenetic neighbourhood. To test this, we combined correlation analyses with systematic linear model grids evaluated by leave-one-out cross-validation and collinearity screening to identify robust, interpretable pre-and post-invasion predictors of ARB invasion success in river biofilm communities.

## Materials and Methods

### River biofilm collection and flume invasion experiments

To compare invasion resistance across natural freshwater biofilms maintained under controlled conditions, river biofilms were collected and transferred into standardised flume systems (Supplementary Figure 1) [17, 32–34]. River biofilms were collected from 20 sites across 12 rivers in 6 European countries (Supplementary Table 1) using artificial exposure units (AEUs), as previously described [17, 32]. Sites were selected to represent contrasting biofilm diversity based on a previous survey [15]. Glass slides were exposed in rivers for 3–4 weeks to allow natural biofilm colonisation. Colonised AEUs were transferred into the laboratory recirculating flume systems containing sterile-filtered river water and acclimated for one week at 20 °C in the dark before the invasion experiments started. Duplicate flumes were then challenged with a single pulse of a model antibiotic-resistant *E. coli* strain at a final concentration of 10^7^ cells mL^-1^ in the recirculating water. Control flumes without *E. coli* addition were run in parallel. Biofilms were destructively sampled from replicate glass slides at Ti (initial biofilm after river colonization), T0 (biofilm after 1 week acclimation immediately before *E. coli* invasion), and during the invasion experiment after 1, 2 (or 3), 7, and 14 days (Tday) (Supplementary Table 2). Biofilm biomass was scraped off the glass slides and DNA was extracted using the DNeasy PowerSoil Pro kit. Further details are provided in Supplementary Methods.

### Invader quantification

To quantify persistence of the invading ARB in total biofilm biomass, invader and total bacterial abundance were monitored. The chosen invader was *E. coli* CM2372, derived from MG1655 Δ*lacZY* [35], carrying the broad-host-range IncP-1α plasmid pG527 encoding kanamycin resistance [17]. Invader persistence within the biofilm communities was quantified by qPCR targeting specific genomic scars of both *E. coli* and pG527, and normalised to total bacterial 16S rRNA gene abundance. Primer sequences, standards, qPCR conditions, and control procedures are provided in Supplementary Methods.

### Amplicon sequencing

To characterise ecological community structure associated with invasion resistance, biofilm communities were analysed using 16S rRNA gene sequencing and phylogeny-derived ecological metrics. Biofilm samples were analysed by 16S rRNA gene amplicon sequencing of the V3–V4 region. Sequences were processed in QIIME2 [36] using DADA2 to generate amplicon sequence variants (ASVs) [37].

### Ecological descriptors of invasion

Shannon diversity, Chao1 richness, and Pielou evenness were calculated from rarefied data for each timepoint (Supplementary Table 4), and Bray-Curtis dissimilarity was computed from the ASV relative abundance table for beta-diversity analyses using PCoA. Additional invader-centred descriptors included abundance-weighted mean phylogenetic distance between all resident ASVs and *E. coli* (MeanDist), relative abundance of ASVs within phylogenetic distance <0.15 to *E. coli* (closeFrac_0.15), Shannon diversity within this close-neighbour subset (Shannon_close_0.15), and changes in Shannon diversity between different timepoints (ΔShannon of Ti→T_0_, T_i_→T_14_, or T_0_→T_14_) (all Supplementary Table 5). Ecological descriptors were classified as predictive descriptors, measurable before or at the time of invasion (e.g. Shannon diversity, phylogenetic neighbourhood occupancy at T_i_ or T_0_ as the closeFrac_0.15), and explanatory descriptors, which describe community restructuring during or after the experiment (e.g. ΔShannon and T14 metrics).

### Statistical analyses, predictive and explanatory ecological modelling of invasion resistance

To identify robust ecological predictors and explanatory descriptors of invasion resistance, both individual correlations and systematic predictive modelling approaches were applied. Invader loss rates were calculated as exponential decay slopes of qPCR-normalised *E. coli* chromosomal marker or plasmid marker abundance over time. Associations between ecological predictors and invader loss were assessed using Spearman rank correlations with *E. coli* chromosomal marker loss rates. Predictive models were fitted independently for T_i_, T_0_, and T_14_ as linear models:

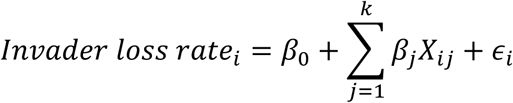

where Invader loss rate_i_ is the invader exponential decay slope in flume i, X_ij_ are the standardised ecological predictors, k is the number of predictors included in the model, and Є_i_ is the residual error. For each timepoint, systematic model grids were generated from the relevant ecological descriptor set containing predictors and explanatory variables for resident biodiversity (Shannon index), occupation of the invader’s phylogenetic neighbourhood (MeanDist, closeFrac_0.15, Shannon_close_0.15) and community stabilisation during transfer into the flume system (either ΔShannon T_i_→T_0_, ΔShannon of T_i_→T_14_ or both, depending on the timepoint). Models were ranked by leave-one-out cross-validation R^2^ (LOOCV R^2^), with root mean squared error (RMSE) and mean absolute error (MAE) reported as complementary error metrics. Models with variance inflation factor (VIF) >5 were excluded, and T_i_ models containing both ΔShannon T_i_→T_0_ and ΔShannon T_i_→T_14_ were omitted to avoid redundant restructuring predictors. Relative predictor contribution was calculated as *β_j_* / ∑|*β_j_*| from standardised model coefficients.

To further explore the overall response in community composition between flumes depending on their invader loss rate, SIMPER (Similarity Percentage) tests were used to identify major taxa contributing to those differences, followed by a PERMANOVA. The flumes within the 25% top quartile of highest invader loss (i.e., lowest invader persistence) were compared to the bottom 25% quartile represented by the flumes with the lowest invader loss (i.e., higher invader persistence). The average dissimilarity, based on Bray-Curtis, was compared between the two sets of flumes both at the beginning of the incubation (T_0_) and at the end (T1_4_). The analysis evaluates whether the centroids of the groups (T_0_ and T_14_) are significantly different in multivariate space by comparing the observed between-group variance to a null distribution generated through permutations of sample labels.

## Results and Discussion

### Natural biofilms exhibiting contrasting ecological structures

To test whether naturally assembled freshwater biofilms provide a sufficiently broad range of biodiversity in the microbial community for exploring the drivers of a model bacterial invasion, we first characterised the diversity and composition of the river-derived communities grown on glass slides before transfer into the flume system. The 41 flumes were established from glass-slide river biofilms grown at 20 sites across 12 rivers in 6 European countries. These biofilms exhibited substantial differences in bacterial community structure at the initial river-derived timepoint (Ti). The 41 selected biofilms covered a broad alpha-diversity gradient, with Shannon index spanning from 5.93 to 9.126, Chao1 richness of 531 ± 191 ASVs, and Pielou evenness of 0.9 ± 0.03 all varying strongly among sites (Figure 1a). This variation occurred mainly among rivers (ANOVA, p-value < 0.05) but also within individual river systems, with pronounced within-river differences observed for rivers like the Glatt (Switzerland), Hirschbach (Germany), and Rouge-Rupt (France) rivers (Figure 1a).

**Figure 1.**
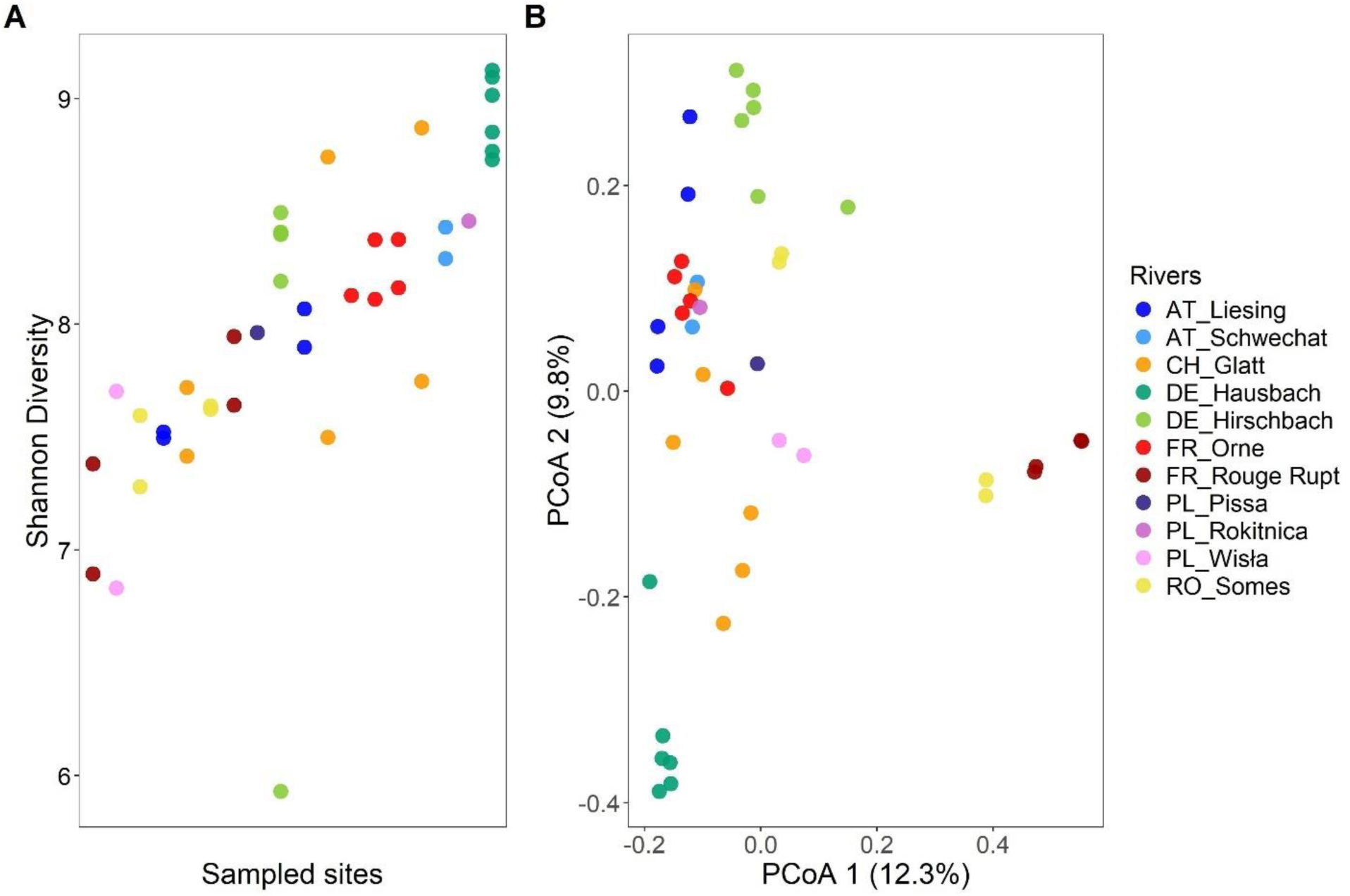
Diversity of the natural biofilms collected in rivers. (A) Shannon diversity index of biofilm communities sampled from the glass slides just after the recovery of biofilms established in AEUs deployed in rivers for 3-4 weeks (T_i_ = time initial). Each point was color labelled according to the river of origin and organized according to the mean Shannon index for the site. (B) Comparison of initial biofilm diversity between sites (beta diversity). The Principal Coordinates Analysis (PCoA) reflects the Bray-Curtis distance of ASV relative abundance of biofilm communities sampled from the glass slides.

Community composition also differed strongly among T_i_ biofilms, with major composition of phyla Pseudomonadota, Bacteroidota and Actinomycetota (Supplementary Figure 2). Principal coordinates analysis based on Bray-Curtis dissimilarity separated biofilms at Ti according to river identity and site-specific community structure, with PCoA1 and PCoA2 explaining 12.3% and 9.8% of the variation, respectively (Figure 1b). Together, these axes explained less than 25% of total variation, indicating high compositional diversity among biofilms originating from different rivers. A small increase in axis explanation was observed for microbial composition throughout time (Supplementary Figure 3). Biofilms from the same river often grouped closer together than biofilms from different rivers, with 56% of the total microbial community composition was driven by the rivers grouping at Ti (PERMANOVA, R^2^ = 0.56, p<0.001). Nevertheless, substantial within-river variation was also apparent, with the highest significant beta dispersion for rivers Glatt (average internal distance = 0.476), Somes (0.467) and Orne (0.455).

Glass-slide biofilms showed comparable dominant taxa and diversity ranges with natural epilithic biofilms collected from the same sites two years prior [15] (Supplementary Figure 4a). Glass biofilms had Shannon indices ranging from 5.93 to 9.12, whereas rock-associated epilithic biofilms ranged from 5.08 to 9.23. Shared ASVs represented 61% for epilthic and 69% for glass samples of total relative abundance across both substrate types for all sites comprised (Supplementary Figure 4b). Thus, although substrate type influenced community composition, the artificial exposure units captured biofilms with diversity levels comparable to natural epilithic communities.

Together, these results show that the invasion experiments were initiated with a broad set of naturally assembled freshwater biofilm communities. This ecological heterogeneity provided the necessary predictor space to test whether resident community structure determines the persistence of an invading antibiotic-resistant bacterium.

### River biofilms differ strongly in permissiveness to ARB invasion

Because invasion ecology theory predicts that resident community structure can constrain establishment of invading organisms through mechanisms such as niche saturation, resource competition, and antagonistic interactions [16, 19, 20, 23, 24], we next tested whether the strong range of biodiversity observed among river biofilms translated into different invasion outcomes for the model antibiotic-resistant *E. coli* strain. Previous work in both macroecology and microbial systems has suggested that invasion resistance emerges from resident community properties rather than the sole propagule pressure alone [20–22]. We therefore assessed whether biofilms differing in diversity and composition also differed in their permissiveness to a model ARB persistence under otherwise standardised invasion conditions.

Following inoculation, both chromosomal and plasmid targets of the invading *E. coli* were quantified by qPCR in all invaded flumes after 24 h, confirming successful mass transfer of the invader from the circulating water phase onto the river biofilms (Figure 2a, Supplementary Figure 5). Despite identical inoculation procedures and similar initial propagule pressure, invader persistence trajectories after 14 days differed strongly among biofilms, ranging from near-complete invader loss to its steady persistence over the incubation time. Invader decline approximately followed first-order decay kinetics, allowing loss rates to be quantified as log decline per day using exponential fitting. Particularly resistant biofilms, such as FR_Or_1, AT_Li_1 and CH_Gl_3, showed rapid invader decline with an average invader loss of 0.32 log of bacterial invader loss per day. On the other hand, other biofilms, including DE_Ha_1, DE_Hi_2 and FR_Ro_2, remained substantially more permissive, presenting an invader loss 10 times lower than the resistant flumes (loss of 0.02 log per day) over the course of the experiment (Figure 2b; Supplementary Figure 5). Replicate flumes exhibited similar invasion trajectories with coefficient of variation around 28%, indicating that invasion outcomes were reproducible and linked to biofilm identity rather than stochastic experimental variation.

**Figure 2:**
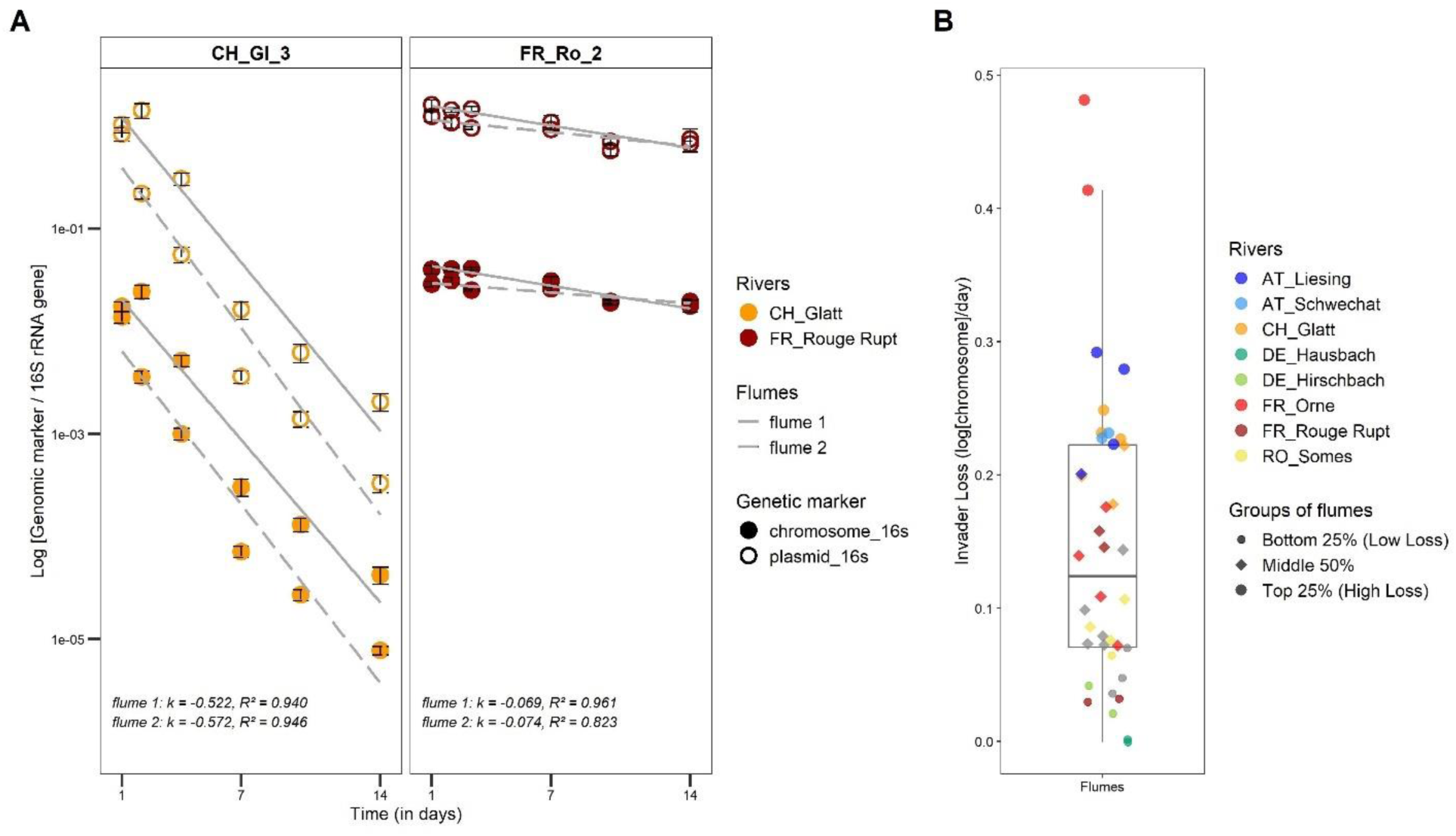
Invasion dynamics of *E. coli* hosting the pG527 plasmid in flume biofilms. (A) Typical invaders loss dynamics in example rivers over time. Fourteen days-decay kinetics of chromosomal (filled markers) and plasmid (empty circles) markers of the invading *Escherichia coli* after normalization to the 16S rRNA gene. Each point represents the average quantification of three qPCR technical replicates with bars representing the standard deviations. Loss rate per day was calculated using exponential decay kinetics from the slope (k). The full dataset of all flumes and locations is given in Supplementary Figure 5. (B) Distribution of invader loss rates. Each point was labelled according to the river of origin.

Because the invader carried the broad-host-range pG527 plasmid, separate qPCR quantification of the invader chromosome and plasmid allowed us to distinguish persistence of the invading host from potential plasmid dissemination into resident biofilm bacteria, as previously applied in complex environmental matrices [11]. Chromosomal and plasmid markers exhibited highly similar decay trajectories, with parallel slopes, indicating that both targets were lost concomitantly. Previous work indicated that, among other, the dissemination of plasmid in natural communities was strongly influenced by the abundance of donor bacteria and its persistence [11]. Here, plasmid transfer was not detected during the course of the biofilm invasion experiments. In this respect, it should be considered that in environmental communities, plasmid transfers occur at a relatively low frequency while transconjugants only persist transiently for fitness reason [11, 38]. Still, plasmids have been shown to persist for decades in the environment, only showing up occasionally in different geographical locations, thus indicating plasmid circulation below detection level [39].

In this set of experiments, we concluded there was weak evidence for plasmid persistence independent of the invading host based on the limited divergence between plasmid and chromosomal decay. The persistence of the introduced AMR determinant was therefore primarily explained by persistence of the invading *E. coli* strain itself, rather than an independent diffusion and establishment of the broad-host-range plasmid in the resident biofilm community, at least over the limit of detection. Based on this result, all subsequent analyses focus on *E. coli* persistence measured by the chromosomal marker. As the relevant measure of ARB invasion success, we use the invader loss rate (*IR* = 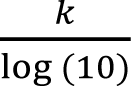 , to obtain positive base-10 values of the chromosomal decay per day, which is inversely proportional to the slope of the first-order decay constant measured (k).

Together, these results confirmed our initial hypothesis that the persistence of our model ARB in freshwater biofilms differ markedly from one community to another, following a fate consistent with ecological theory predicting that resident community properties influence invasion outcomes [19, 20, 22]. Next we tested whether this variation could be predicted from three ecological dimensions expected to influence biotic resistance: i) resident community diversity as more diverse communities may occupy available niche space more effectively [16, 19, 23]; ii) community restructuring during transfer into the flume system, because diversity loss may indicate disturbance-associated niche opening [27, 29, 30]; and iii) invader-specific niche occupancy, because communities containing more abundant or diverse close phylogenetic neighbours of *E. coli*, or a higher average similarity to the invader, may impose stronger competition for overlapping ecological space [40–42].

### Community diversity predicts invasion resistance

Biotic resistance theory predicts that diverse resident communities should be less permissive to invasion because they more completely occupy available niche space, deplete limiting resources, and increase the likelihood of competitive or antagonistic interactions [19, 20, 26]. We therefore tested whether natural variation in biofilm diversity across the pan-European flume experiment explained the heterogeneity of *E. coli* invasion success under standardised invasion conditions.

Community structures were compared at T_i_, T_0_, and T_14_ for each flume. Biofilm diversity changed moderately during transfer and acclimation in the flume system (and after the invasion experiment started). Alpha diversity based on the Shannon index did not change significantly over time when all flumes were analysed together (*P*=0.226, ANOVA) (Supplementary Figure 6). Individual flumes nevertheless showed significant gains or losses in diversity, with Shannon index decreasing by approximately 10% on average (*P*=0.00878). Strong diversity losses were restricted to a few river systems, including the Pissa (Poland), Hausbach (Germany), and Hirschbach (Germany) rivers, where Chao1 richness declined by more than 50% (Supplementary Figure 6). Control flumes without invader addition showed similar diversity trajectories, indicating that these shifts reflected acclimation to the flume system rather than a specific effect of the *E. coli* invasion (*P*=0.5, ANOVA, Supplementary Figure 7). Biofilm bacterial biomass, estimated from 16S rRNA gene abundance, remained broadly stable throughout the experiment, with no indication of biomass collapse over time (*P*=0.6, ANOVA, Supplementary Figure 8).

Resident biofilm diversity during invasion strongly explained *E. coli* loss. Shannon diversity was weakly and non-significantly correlated with invasion loss rate at T_i_ (Spearman *R*=-0.037, *P*=0.85), but showed strong negative correlations at T_0_ (*R*=0.52, *P*<0.001) and T_14_ (*R*=0.63, *P*<0.001) (Figure 3A). These correlations show that more diverse biofilms during the invasion phase were more resistant to the *E. coli* invader persistence (higher invader loss). Evenness showed the same temporal pattern, with weak, non-significant association at T_i_ (R*=*0.12, *P*=0.53), a moderate association at T_0_ (R=0.36, *P*=0.038), and a strong association at T_14_ (R=0.57, *P*<0.001) (Figure 3B). Chao1 richness was not associated with invader loss at T_i_ (R=-0.13, *P*=0.5), but became predictive pre-invasion at T_0_ (R=0.41, *P*=0.016) and explanatory post-invasion at T_14_ (R=0.55, *P*<0.001) (Figure 3C).

**Figure 3:**
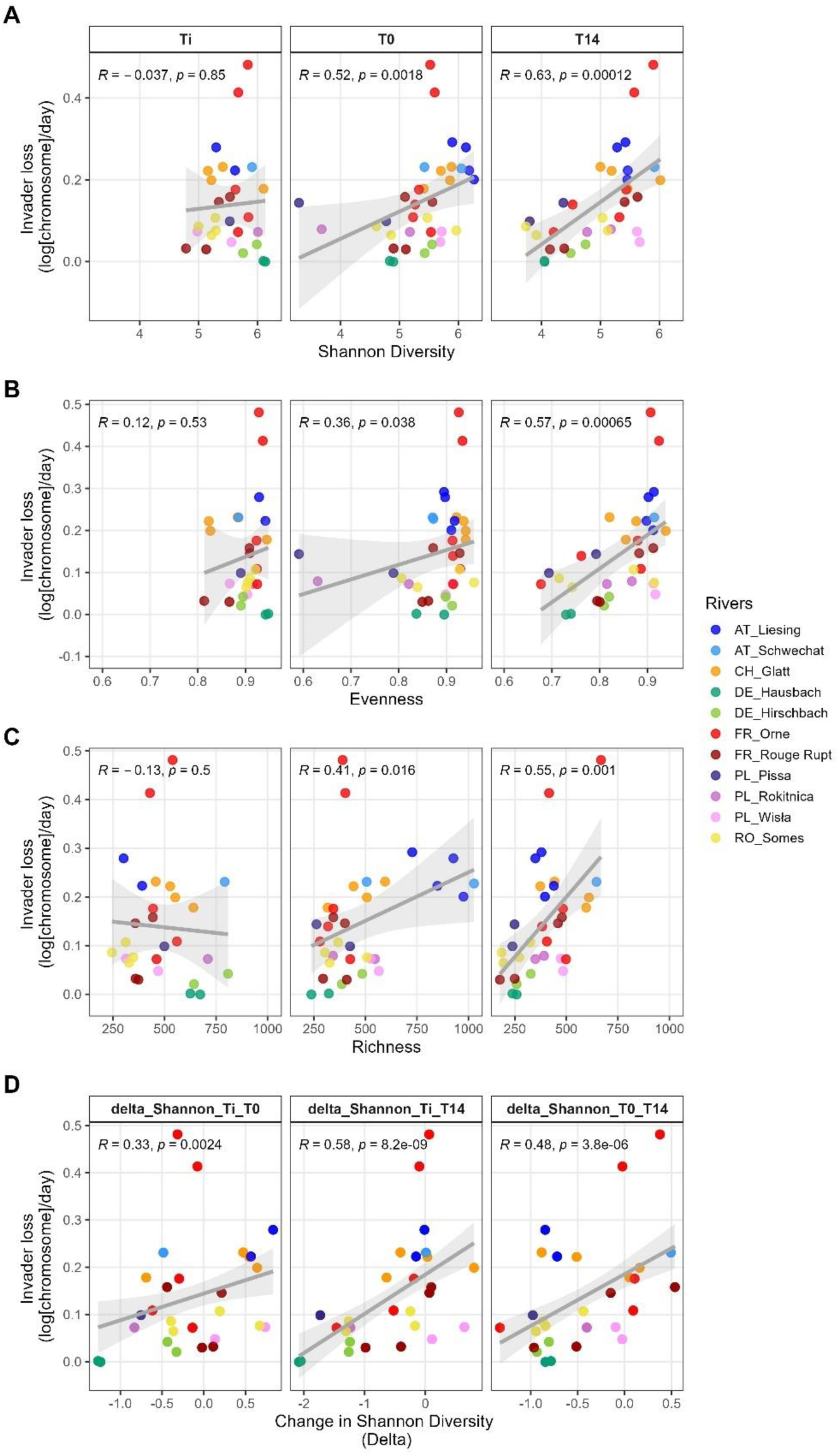
Correlation of diversity metrics of the biofilms with invader loss rates at different timepoints. Correlation of invader loss rate based on Spearman rank correlation with (A) Shannon diversity at T_i_, T_0_, T_14_, (B) Pielou Evenness at T_i_, T_0_, T_14_, (C) Chao1 Richness at T_i_, T_0_, T_14_, (D) Diversity loss (ΔShannon) for the transitions from T_i→_T_0_, T_i_→T_14,_ T_0_→T_14_.

These results support the diversity-invasion effect in natural river biofilms, while adding an important temporal qualification. Whether the diversity has changed because of the transition from river-to-flume, or because loosely interacting bacteria were lost from the biofilm during the experiment, the fact is that microbial community has changed, and the biodiversity characteristics of the mature biofilm was associated with its permissiveness with respect with the invader persistence. Furthermore, diversity-based invasion resistance is generally interpreted within the framework of niche saturation and competitive exclusion, where more diverse resident communities reduce ecological opportunity for invaders through more complete occupation of available niche space [19, 20, 26]. In structured environmental microbiomes, high microbial diversity was previously associated with reduced ARG accumulation in soils, whereas comparable relationships were not observed in more dynamic riverbed systems in the long term, suggesting that environmental stability may modulate diversity-based barrier effects towards invasion [15]. The present experiment extends these observations by demonstrating that natural ranges of biodiversity among dynamic river biofilms are sufficient to predict ARB persistence well if biofilms are kept under stable conditions. During the course of the flume experiments, an invasion test was run in parallel using an alternative invader, *Enterococcus faecium* Com12Δ*sodA* hosting the pSF3 conjugative plasmid, on biofilms sampled in the Orne river. The *E. faecium* invader disappeared at the same pace as its *E. coli* counterpart (Supplementary Fig. 9), indicating that diversity-associated permissiveness of biofilm community likely dominates the fate of invading bacteria over the identity of the invading bacteria itself, although this conclusion would need to be substantiated with further experiments for generalization.

The comparatively weak diversity-invader persistence relationship at T_i_ further indicates that invasion resistance is not simply determined by the original level of diversity, but rather by the ecological structure after stabilisation before (T_0_) and after invasion (T_14_). Previous work was unable to determine a diversity-based barrier effect from riverbed microbiomes [15].

Here we have shown that higher Shannon diversity, which integrates both richness and evenness, was consistently the strongest individual diversity predictor, suggesting that both the number of resident taxa and their relative abundance distribution contribute to invasion resistance. The stronger relationships at T_0_ and T_14_ compared with T_i_ indicated that invasion resistance depends not only on diversity itself, but also on whether this diversity is retained during the transition from the river and over time in the flume system. We next tested whether the degree of community restructuring during transfer and acclimation, expressed as the loss in Shannon diversity over time (ΔShannon) between the different timepoints, which also explained variation in invasion resistance.

### Community restructuring during ecosystem transition shapes invasion resistance

Because diversity measured after acclimation predicted invasion resistance more strongly than diversity at river collection, we next tested whether diversity retention during the river-to-flume transition and during the flume experiment itself explained invasion outcomes retrospectively.

Biofilms differed strongly in the way they retained diversity following transfer into the flume system. Some communities remained comparatively stable throughout acclimation and invasion, whereas others underwent substantial diversity loss despite similar initial diversity values (Figure 3D). The loss in diversity expressed as ΔShannon calculated across the full transition from T_i_ to T_14_ represented the strongest explanatory descriptor for stabilisation and was significantly associated with invasion loss (Spearman *R*=0.58, *P*<0.001). ΔShannon T_0_→T_14_ showed a lower but statistically significant correlation with invader loss (*R*=0.48, *P*<0.001). Lastly, ΔShannon T_i_→T_0_ , as the change of the community observed after one week flume acclimation, correlated the least of invasion outcomes (*R*=0.33, *P*<0.001). These results show that communities retaining Shannon diversity during acclimation and maturation were more resistant to the *E. coli* persistence (higher invader loss), whereas communities undergoing stronger diversity loss became more permissive (lower invader loss).

These findings suggest that invasion resistance is shaped not only by the diversity level reached in the mature biofilm, but also by the degree of community restructuring during ecosystem transition. Communities that retained diversity may have preserved niche occupancy and interaction structure, whereas diversity loss may reflect disturbance-associated niche opening that increased ecological opportunity for the invader [27, 29]. Thus, ΔShannon provides a simple proxy for biofilm stabilisation during transfer and helps explain why Ti diversity alone was only weakly predictive of invasion resistance.

### Occupation of the invader’s phylogenetic neighbourhood predicts invasion resistance

Although community diversity and its stability explained a substantial proportion of variation in invasion resistance, diversity metrics alone do not capture whether resident taxa occupy ecological niches similar to that of the invader. Ecological and evolutionary theory predicts that invasion success should depend particularly on the overlap of the ecological space between the invader and resident competitors, a concept often described as phylogenetic limiting similarity or Darwin’s naturalisation hypothesis [40–42]. Because phylogenetic relatedness can partially reflect shared metabolic capabilities, resource requirements, and environmental preferences in microbial systems [43], we next tested whether the phylogenetic relatedness (based on 16S rRNA gene) of the community structure with the invader, explained invasion resistance beyond total diversity alone.

Mean phylogenetic distance between the resident community and *E. coli* (MeanDist) was only weakly associated with invasion resistance and did not represent a significant individual predictor at any timepoint (T_i_: Spearman *R*=0.17, *P*=0.4; T_0_: *R*=-0.24, *P*=0.16; T_14_: *R*=-0.24, *P*=0.18, Figure 4a). Thus, average community-wide phylogenetic similarity to the invader alone did not explain ARB persistence. In contrast, metrics specifically describing the invader’s local phylogenetic neighbourhood showed substantially stronger relationships with invasion outcomes. The relative abundance of ASVs within a phylogenetic distance threshold of <0.15 to *E. coli* (closeFrac_0.15) was positively associated with invader loss at T_0_ (*R*=0.38, *P*=0.026) and T_14_ (*R*=0.41, *P*=0.018), whereas no relationship was observed at T_i_ (*R*=0.012, *P*=0.95) (Figure 4). Likewise, Shannon diversity within this close phylogenetic neighbourhood (Shannon_close_0.15) was positively associated with invasion resistance at T_0_ (*R*=0.36, *P*=0.037) and particularly strongly at T_14_ (*R*=0.55, *P*<0.001), while T_i_ again showed no significant relationship (*R*=0.12, *P*=0.54) (Figure 4). Therefore, the results indicated that biofilms containing more abundant and more diverse communities with close phylogenetic relationship to *E. coli* were substantially more resistant to invasion.

**Figure 4:**
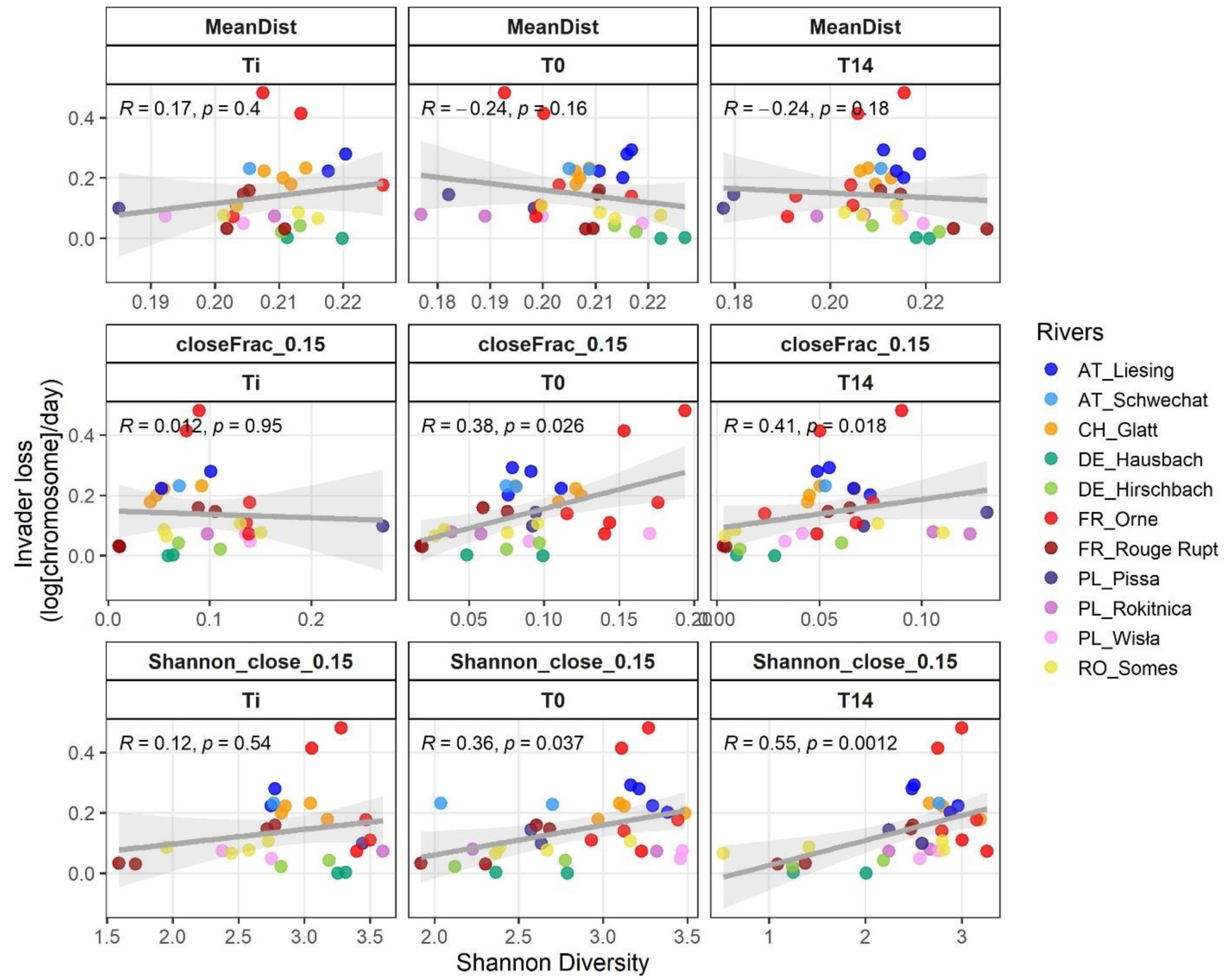
Correlation of invader-centred phylogenetic neighbourhood metrics with invader loss rates. Invader loss rates were correlated with ecological metrics describing the phylogenetic relationship between resident biofilm communities and the invading *E. coli* strain using Spearman rank correlations. (A) MeanDist represents the abundance-weighted mean phylogenetic distance of all resident ASVs to *E. coli*. (B) closeFrac_0.15 represents the summed relative abundance of ASVs within a phylogenetic distance <0.15 from *E. coli*. (C) Shannon_close_0.15 represents Shannon diversity calculated only within this close phylogenetic neighbourhood. Correlations are shown separately for T_i_, T_0_, and T_14_.

These findings suggest that invasion resistance was governed only weakly by community-wide phylogenetic relatedness to the invader, likely too coarse to capture the competitive interactions most relevant for ARB persistence. Instead, invasion resistance increased when communities contained larger and more diverse pools of taxa closely related to the invader, consistent with competitive exclusion by organisms occupying overlapping ecological space [40, 41]. Similar patterns have previously been observed in microbial invasion experiments, where phylogenetically or functionally similar resident taxa reduced the establishment success of invading bacteria [26, 42]. Importantly, phylogenetic similarity should be interpreted cautiously here as a proxy for potential ecological overlap rather than direct functional equivalence. Nevertheless, the stronger predictive performance of invader-neighbourhood metrics compared with average community distance supports the idea that biotic resistance emerges not only from overall diversity, but also from whether the invader’s specific niche space is already occupied within the resident community.

### Integrating ecological descriptors improves prediction and explanation of invasion resistance

After testing diversity, community restructuring, and invader-specific phylogenetic neighbourhood structure separately, we next asked how these ecological dimensions combine to either predict invasion resistance before or during the invasion process or explain observed invasion success retroactively. For each timepoint, we generated all linear models from the final ecological descriptor set, ranked them by leave-one-out cross-validated performance, and screened them for collinearity using VIF, excluding models with VIF>5. This allowed us to compare predictive strength across models while retaining only interpretable predictor combinations.

At T_0_ and T_14_, the best-performing models consistently combined Shannon diversity with mean phylogenetic distance to *E. coli* (Figure 5). At T_0_, the Shannon + MeanDist model achieved a LOOCV R^2^ of 0.241, clearly outperforming Shannon alone as the best individual descriptor (LOOCV R^2^=0.077; Figure 5, Supplementary Figure 10). The seven best T_0_ models all contained both Shannon and MeanDist, but adding further descriptors to this simplest two-variable model reduced cross-validated performance. Standardised model coefficients indicated that Shannon was the strongest contributor at T_0_ (β=-0.161), followed by MeanDist (β=0.121). Thus, diversity was the dominant factor in predicting invasion before it happened, but phylogenetic structure added clear complementary information.

**Figure 5:**
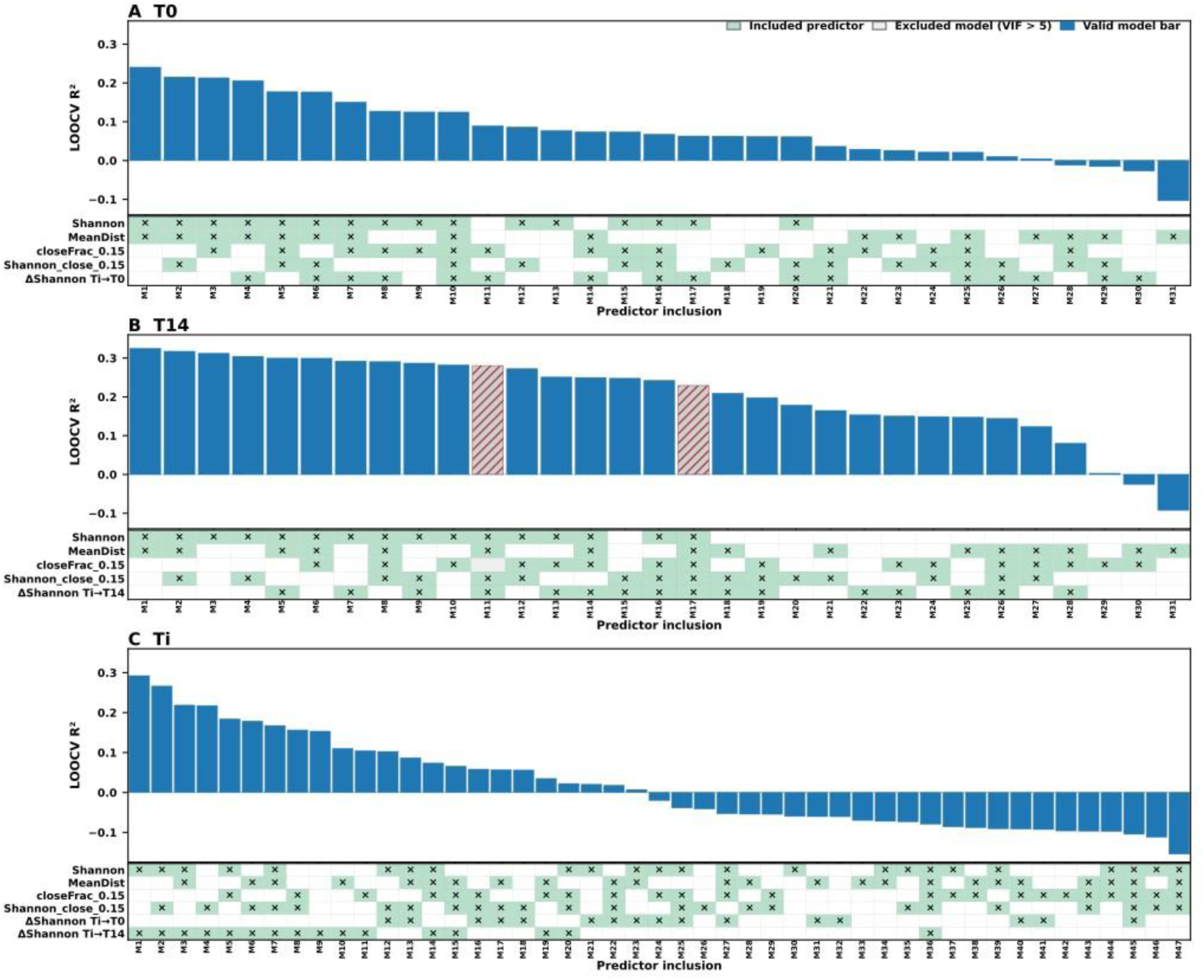
Predictive and explanatory ecological modelling of ARB invasion resistance across river biofilms. Systematic linear model grids were generated independently for (A) T_0_, (B) T_14_, and (C) T_i_ to identify ecological descriptors of invader loss rates. Models were ranked by leave-one-out cross-validation R² (LOOCV R²). Horizontal bars indicate model predictive performance, while the descriptor matrix indicates inclusion of ecological descriptors within each model. Colours in the descriptor matrix indicate whether descriptors were retained or excluded following variance inflation factor (VIF) screening. Descriptor sets included Shannon diversity, MeanDist, closeFrac_0.15, Shannon_close_0.15, and ΔShannon restructuring metrics. For T_i_, models containing both ΔShannon T_i_→T_0_ and ΔShannon T_i_→T_14_ were excluded to avoid redundant restructuring descriptors.

At T_14_, Shannon + MeanDist again ranked as the best model (LOOCV R^2^=0.325) explaining invasion success after invasion happened, but Shannon alone as the best individual predictor performed nearly as well (LOOCV R^2^=0.312; Figure 5b, Supplementary Figure 10). Component strength also shifted strongly towards Shannon, with a much larger standardised coefficient for Shannon (β=-0.157) than for MeanDist (β=0.038). This indicates that once biofilms had matured, traditional diversity captured most of the predictable variation in invasion resistance, whereas MeanDist contributed only marginal additional information.

The T_i_ model space displayed a different pattern (Figure 5c). Models based only on static T_i_ community properties performed weakly in predicting subsequent invasion success, whereas the best Ti model combined Shannon diversity with ΔShannon T_i_→T_14_ (LOOCV R^2^=0.292), thus only retroactively explaining observed invasion dynamics in our experimental system. All top-performing Ti models contained diversity loss during transition, indicating that invasion resistance of the original river-derived communities could only be explained retrospectively when subsequent community restructuring was included. In this model, ΔShannon was the strongest component (β=-0.172), with Shannon as the main co-descriptor (β=-0.110), showing that diversity retention during transition was more informative than initial diversity alone. Nevertheless, these dynamics might be specific to an experimental system where the change in diversity can be observed and tracked, and this change can be used as an explanatory variable.

Explaining approximately one-third of the variation in invasion resistance from only a small number of community-level descriptors represents substantial predictive power for a natural ecological system. Given the large number of unmeasured environmental variables, stochastic demographic processes, and complex microbial interactions influencing community dynamics, perfect prediction is neither expected nor ecologically realistic [44–46]. The observed predictive performance therefore suggests that biodiversity, community restructuring, and invader-specific niche occupancy capture fundamental ecological processes governing ARB establishment.

Together, these results show that ARB invasion resistance is quantitatively predictable from ecological community properties, but that predictor importance depends on biofilm state. In resident river communities, prediction depends primarily on whether communities retain diversity during transition into the experimental ecosystem. In acclimated biofilms, invasion resistance is best predicted by combining diversity with phylogenetic structure. In mature biofilms, Shannon diversity alone captures most of the predictable resistance signal. This supports a model in which freshwater biofilm invasion resistance is not stochastic, but emerges from the interaction between biodiversity, community stabilisation, and invader-specific niche structure.

### Distinct ecological guilds characterize resistant and permissive biofilms

Having identified diversity, community restructuring, and invader-specific niche occupancy as major predictors of invasion resistance, we finally asked whether resistant and permissive biofilms were consistently associated with distinct taxa or guilds. Rather than interpreting individual taxa as direct causal drivers, the SIMPER analysis presented here aimed to identify community-level ecological signatures associated with contrasting invasion outcomes.

SIMPER analyses revealed consistent compositional differences between highly invasion resistant (top 25% flumes with highest invader loss) and highly permissive biofilms (bottom 25% flumes with lowest invader loss) at both T_0_ and T_14_. A significant PERMANOVA (P=0.003) suggested that the microbial community structure (Bray-Curtis dissimilarity between ASVs relative abundance) differed between the two groups of flumes. The top 30 ASVs discriminating between permissive and resistant biofilms at T_0_ and T_14_ were grouped together by their class. (Figure 6). The ASVs contributing the most for the difference between permissive and resistant, were mostly discriminant at T_14_ when compared to T_0_. This is in agreement with Shannon diversity at T_14_ best explaining invader loss (Figure 3c-4). Furthermore, the right panel portrays the average relative abundance of each class in either group of flumes.

**Figure 6.**
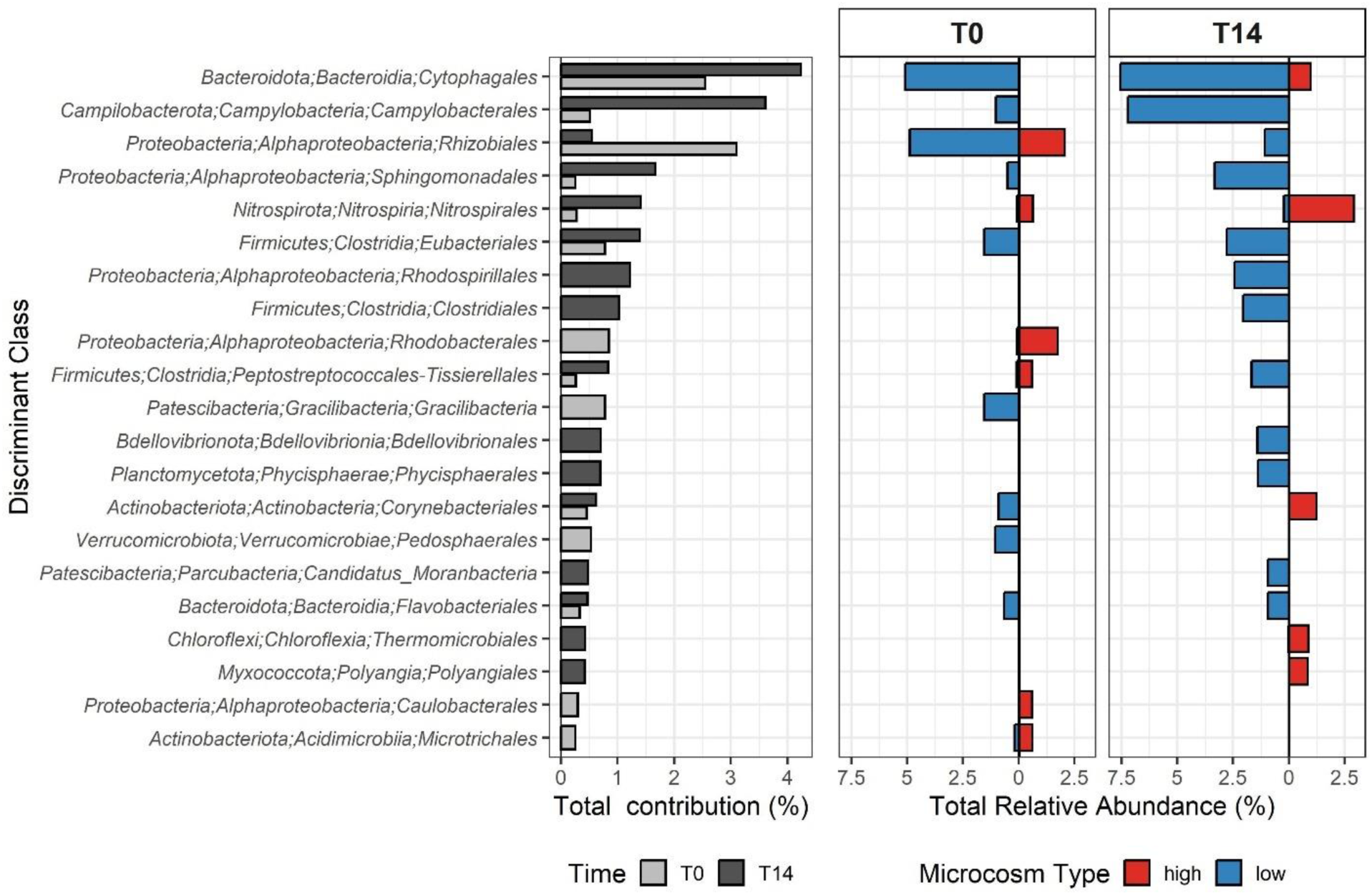
SIMPER analysis to identify discriminant ASVs. ASVs and their associated taxonomy discriminant between flumes with low or high invader loss rate, and at T0 and T14. Microcosms type refers to the flumes “high” – with resistant-to-invasion biofilms with highest invader loss; or “low” – with permissive biofilms with lowest invader loss.

Communities associated with slow *E. coli* invader loss were enriched in several taxa commonly linked to structured biofilms, particle-associated lifestyles, or complex anaerobic and oligotrophic community organisation. The strongest permissive-associated groups included *Cytophagales*, *Campylobacterales*, *Rhizobiales*, *Sphingomonadales*, *Eubacteriales*, *Rhodospirillales*, and *Clostridiales*. *Cytophagales* represented the strongest discriminating group at both T_0_ and T_14_, contributing more than 4% to overall community differentiation (Figure 6). Several of these groups, particularly *Cytophagales*, *Campylobacterales*, and *Rhizobiales*, increased further in relative importance in resistant communities between T_0_ and T_14_, paralleling the increasing predictive strength observed for mature biofilms.

In contrast, invasion-resistant biofilms with rapid *E. coli* loss were associated primarily with *Nitrospirales* and *Rhodobacterales*, particularly at T_14_ (Figure 6). These communities additionally exhibited weaker enrichment of several copiotrophic or metabolically flexible taxa, suggesting broader ecological differences in community organisation between resistant and permissive states.

The SIMPER results therefore suggest that invasion resistance is associated not only with higher diversity, but also with broader ecological community structure. As discussed before, such communities may provide stronger niche occupation and reduced ecological opportunity for invading ARB. In contrast, permissive communities contained higher relative abundances of taxa often associated with metabolically flexible or less spatially structured lifestyles, potentially reflecting communities with greater ecological openness towards invading bacteria. Importantly, these taxa should not necessarily be interpreted as direct determinants of invasion resistance. Instead, they likely represent ecological indicators of broader community states associated with resistant or permissive biofilm organisation.

## Conclusions

Our results demonstrate that environmental persistence of an invading antibiotic-resistant bacterium is not stochastic, but instead emerges from measurable ecological community structure. Across naturally assembled freshwater biofilms, invasion resistance could be quantitatively predicted from three major ecological dimensions: overall biodiversity, community stabilisation during ecosystem transition, and phylogenetic neighbourhood structure relative to the invader.

Among these factors, biofilm overall diversity represented the strongest and most consistent predictor of invasion resistance, while phylogenetic similarity and diversity retention during community restructuring provided important additional predictive or explanatory power depending on biofilm state. These findings support the view that invasion resistance emerges from integrated ecological community properties rather than from individual antagonistic taxa alone. Mainly, we show that stable freshwater microbial communities may function as natural ecological barriers against AMR establishment, but that invader-specific ecological overlap also contributes substantially to invasion success.

These findings align closely with ecological invasion theory, which predicts that invasion success is governed by niche availability, competitive exclusion, and disturbance-mediated restructuring of resident communities [19, 20, 47, 48]. In this framework, mature and diverse biofilms likely reduce ecological opportunity for invading ARB through more complete resource utilisation, stabilised interaction networks, and occupation of invader-relevant niche space. Conversely, community restructuring and diversity loss may transiently increase ecological openness and facilitate establishment of invading resistant bacteria. Our results therefore support the view that microbial invasion resistance in freshwater systems emerges from classical ecological principles operating within complex environmental microbiomes.

More broadly, our results suggest that the environmental risk associated with ARB release depends not only on propagule pressure or ARGs (horizontal) mobility [49], but also on the ecological structure and stability of the receiving microbiome. Integrating ecological community properties into environmental AMR surveillance and risk assessment may therefore improve our ability to predict where resistant bacteria are most likely to persist and establish in natural ecosystems.

## Acknowledgements

We thank Hélène Guilloteau from LCPME for constructing the two strains used in this study.

## Author contributions

Conceptualization of the study and sampling strategy: EC, CM, XB, UK, TUB, GG, HB, SG, AGS, ES, CC, NK, MP, FW, MW; Flumes strategy testing: EC, GG, UK, ES, CC; Identification of national sampling locations, flume operation, sampling, metadata collection & sample processing: UK, GG, EC, XB, ID, EDE, SG, AGS, UO, ER, MS, ES, AT-M, CC, NK, MP, JV, FW, MW, HB, CM; Bioinformatic sequence analysis: EC, AXE; qPCR data analysis: EC, APDM, CM; Predictive modelling: AXE, UK; Data curation and validation: EC, CM; Data interpretation: EC, CM, UK, GG, HB, TUB; Visualization of data: EC, UK, GG; Funding acquisition: UK, CC, NK, MP, FW, MW, HB, CM, TUB; Supervision: UK, CC, NK, MP, JV, MW, HB, CM, TUB; Writing -original draft: EC, UK, CM; Writing review and editing: all authors. All authors have read and approved the final version of the manuscript.

## Data availability

The datasets supporting the conclusions of this article are included within the article and its additional supplementary files or are available through the corresponding author upon reasonable request.

Raw 16S rRNA sequence data are currently being deposited in the NCBI Sequence Read Archive (SRA) and accession numbers will be provided in a revised version of this preprint and prior to peer-reviewed publication.

