## Supplementary Material for "Microbial community diversity predicts invasion resistance of freshwater biofilms against antibiotic-resistant bacteria"

Running title: Predicting ARB Invasion in River Biofilms

Elisa C. P. Catão[1,2,*] , Uli Klümper[3,*], Giulia Gionchetta[4,5,*], Xavier Bellanger[1], Amélie Porteu de la Morandière[1], Alan Xavier Elena[3], Irina Dielacher[6], Sonia Galazka[7], Agata Goryluk-Salmonowicz[8], Elena Radu[6,10], Mateusz Szadziul[8], Edina Szekeres[11], Adela Teban-Man[11], Cristian Coman[11], Norbert Kreuzinger[6], Magdalena Popowska[8], Julia Vierheilig[6,12], Shauna O'Shea Boland[10], Fiona Walsh[9], Markus Woegerbauer[7], Helmut Bürgmann[4], Thomas U. Berendonk[3,], Christophe Merlin[1,#]

### Supplementary Methods

**River biofilm collection and flume setup**

Each AEUs consisted of 9 microscope glass slides mounted on a plexiglass holder attached to a concrete plate covered with a metal plate, allowing tangential water flow along the glass slides through a metal mesh (Suppplementary Figure 1). Platforms containing eight AEUs each were deployed per river site for approximately 3-4 weeks. After colonisation, AEUs were transported to the respective laboratories and installed into recirculating flumes (3 AEUs per flume) containing 8 L of filtered river water. River water was sterile-filtered through 0.45 µm PES membrane filters, recirculated at 100 mL $min^{-1}$, and 10% of water volume was replaced weekly to replenish nutrients. Biofilms were acclimated for one week before invader addition in half of the flumes, with the other half serving as non-invasion controls.

**Additional invader quantification**

To quantify the persistence of an additional bacteria in the flumes, four flumes (in duplicates; in french flumes) were subjected to the introduction of another invader, *Enterococcus faecium* strain (CM2451), a derivative of strain D34 with chromosomal Com12Δ*sodA* and a pSF3 Δ*vanZ* plasmid. This bacterium was cultured in BHI with 10 mg/mL erythromycin, and specific primers were used to monitor both the chromosomal and the plasmid markers as for *E. coli*.

**DNA extraction and qPCR**

For each destructive sampling of biofilms, an individual glass slide with grown biofilm was randomly removed from each of the three replicate AEUs of each flume. The remaining surrounding water was removed, and biofilms were scraped from each individual glass slide inside an individual 50 mL centrifuge tube using a cell scraper. One mL of detergent solution (0.9 % sterile NaCl solution with 0.05 % of Tween80 was added to support the scrapping process and allow complete detachment of the biofilm from the glass slide. Samples were then centrifuged at 13,000 × g for 5 min, the supernatant carefully removed by pipetting and the cell pellet stored at -20 °C. DNA was extracted using the Qiagen DNeasy PowerSoil Pro kit according to the manufacturer's instructions.

qPCR assays targeted an *E. coli* chromosomal marker (MG1655Δ*lacZY* amplicon target), plasmid pG527 (amplicon target overlapping the plasmid backbone and Tn*7* right end (*att*Tn*7*R)), and bacterial 16S rRNA genes (Supplementary Table 3). For French additional flumes, *E. faecium* Com12 Δ*sodA* chromosomal marker (Δ*sodA* amplicon target) and plasmid pSF3 Δ*vanZ* (Δ*vanZ* amplicon target) were monitored as well. As a qPCR standard, the recombinant plasmid pFTs (a pEX-A258; pUC derivative), containing all targets, was extracted from the strain E. coli Top10 (CM2455) using the Wizard plus SV Miniprep DNA Purification System (Promega, Madison, WI, USA) according to the manufacturer instructions and linearized by XbaI (Promega) before being purified with the QIAquick PCR purification kit (Qiagen). Chromosomal and plasmid markers were quantified using TaqMan chemistry, whereas 16S rRNA genes were quantified using SYBR chemistry. Control flumes and pre-invasion samples were used to confirm absence of invader markers. All samples were quantified in technical triplicates.

Reactions were performed in technical triplicates in a qPCR machine different for each partner (Germany: MasterCycler RealPlex; France: StepOne Plus; Switzerland: ; Poland: ;Romania: StepOne Plus. All reactions were performed for a final volume of 20 µL or 25 µL with 1X final concentrations of either qPCR Master Mix for the 16S rRNA gene or 1X Probe qPCR Master Mix for the other targets. Each primer was added at a final concentration of 250 or 300 nM and the probe at a final concentration of 80 nM. All PCR programs consisted of initial denaturation at 95 °C for 10 min and 45 cycles of denaturation (95 °C; 15 s) and annealing and elongation (60 °C; 1 min). Standard curves for either of the targets were created during every qPCR run, using the above-described standard plasmid (containing both targets) with the standard target concentrations ranging from $10^6$ to $10^1$ copies per reaction. Standard curves for the 16S rRNA were created with a ¼ fold dilution starting from $10^7$ to avoid the inhibition of the reaction with a plateau reached between $10^3$ and $10^2$ copies per reaction. Standard curves with amplification efficiency 0.9–1.1 and R2 ≥ 0.99 were accepted and melting curve analysis was performed to assess the amplicons' specificity in the case of 16S rRNA gene quantification. Screening for PCR inhibition was performed by spiking the standard plasmid into the DNA samples. No inhibition was detected in any of the samples. Importantly, no amplification of the *E. coli* target was observed for any of the samples from the control treatment, hence verifying that exclusively the focal *E. coli* strain CM2372, and no unspecific environmental bacteria are detected using this set of primers and probe. The limit of quantification was calculated for each individual qPCR run according to the MIQE guidelines [1]. The absolute abundance of genes was finally expressed as gene copies/cm2 on the glass slides and the relative abundance of *E. coli* CM2372 was calculated as the ratio of target copies per copy of the 16S rRNA gene.

**16S rRNA gene sequencing and processing**

The DNA extracts were sent to the IKMB Kiel University (Germany), and the 16S rRNA genes were amplicon sequenced on an Illumina Novaseq using the primers (v3f: CCTACGGGAGGCAGCAG; v4r: GGACTACHVGGGTWTCTAAT). Reads were processed in QIIME2 version 2021.11 [2] using DADA2 [3]. Paired reads were merged, quality-filtered, and resolved into ASVs. Rare ASVs below 0.1% mean sample depth, mitochondria, and chloroplast sequences were removed after classification against SILVA 138. Alpha diversity was calculated after rarefaction to 3700 reads.

### Supplementary Figures

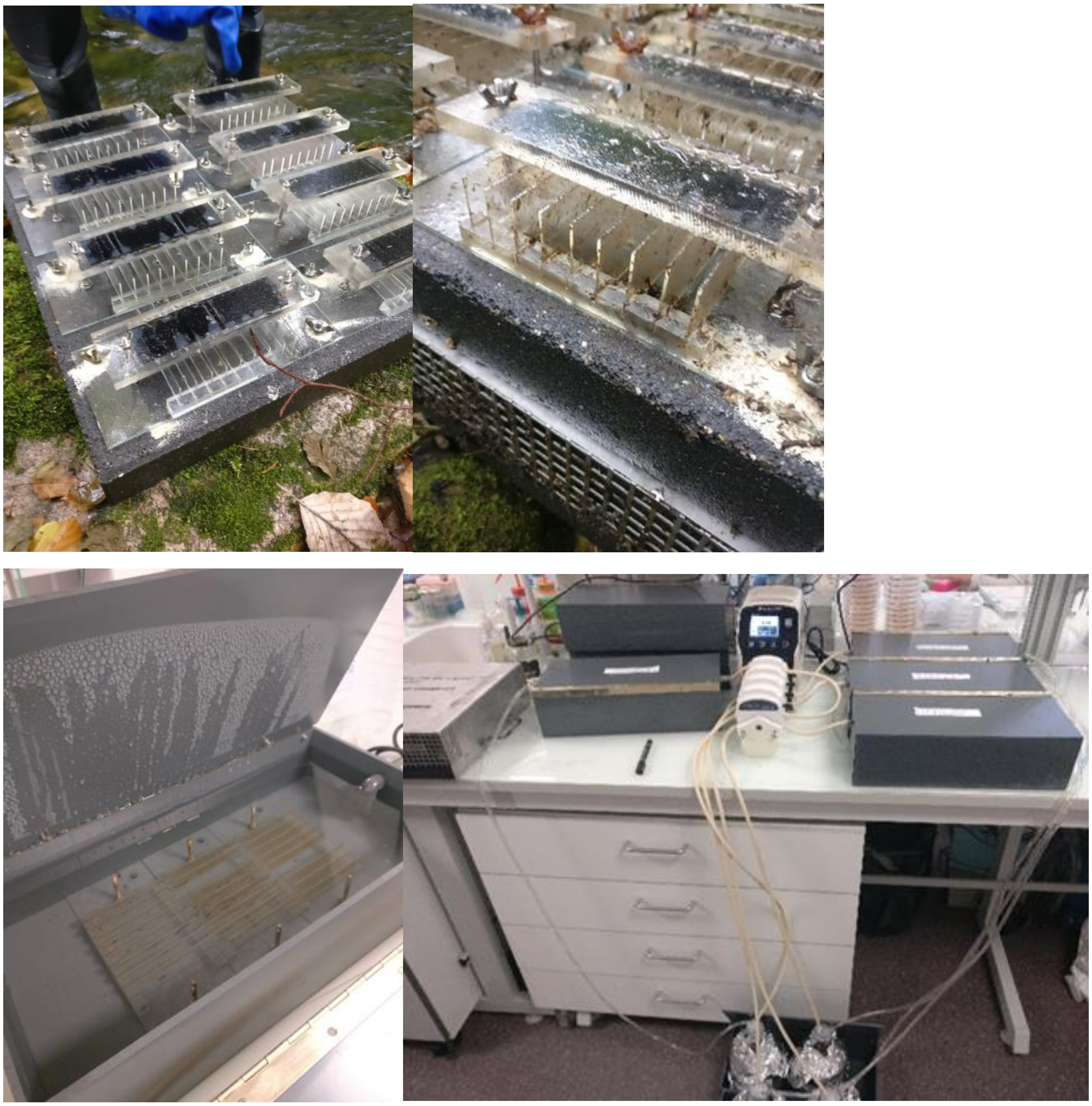

**Supplementary Figure 1.** *In situ* (top) and laboratory (bottom) set-ups for this study. Glass slides were kept in the metallic and concrete cages on the bottom of the rivers for around 1 month for biofilm formation, and the acrylic supports with 9 glass slides each (top) were transferred to the laboratory upon arrival for the closed recirculation of filtered river water on opaque flumes (bottom).

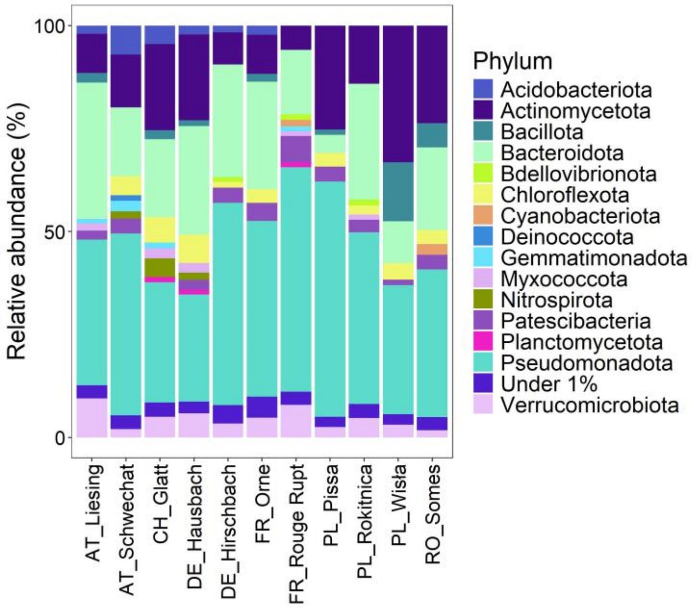


**Supplementary Figure 2.** Bacterial taxonomic community composition at phylum level in all river biofilms grown over the glass slides, sampled just after the 1-month immersion in the river (Ti). Phyla with < 1% relative abundance were grouped in "Under 1%". Each bar corresponds to the sum of all abundances for the replicates obtained at Ti for each river site before performing the relative abundance.

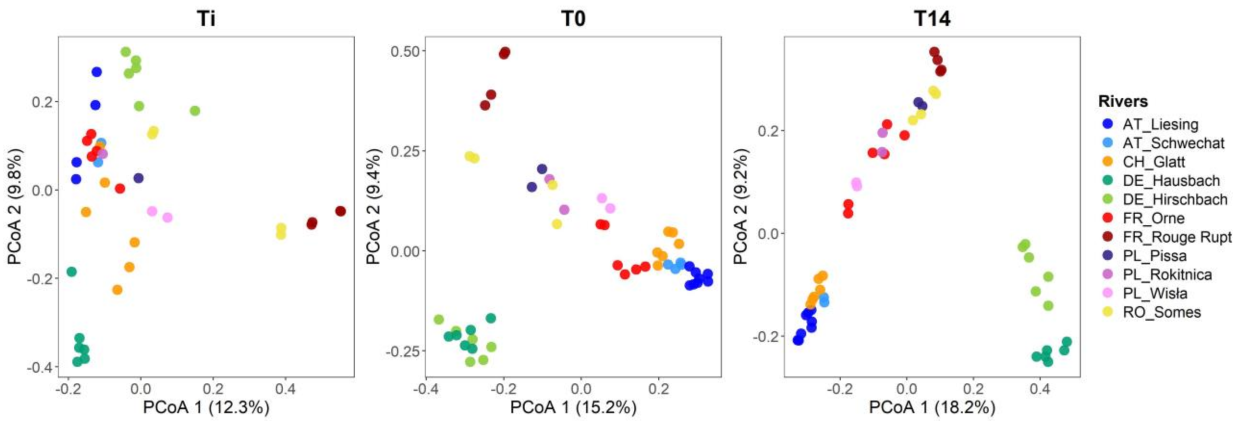


**Supplementary Figure 3.** Beta diversity of the biofilms for the comparison of initial biofilm diversity between sites (Ti) and in the beginning of the flumes (T0) and at the end of the invasion experiment (T14). The Principal Coordinates Analysis (PCoA) reflects the Bray-Curtis distance of ASV relative abundance of biofilm communities sampled from the glass slides.

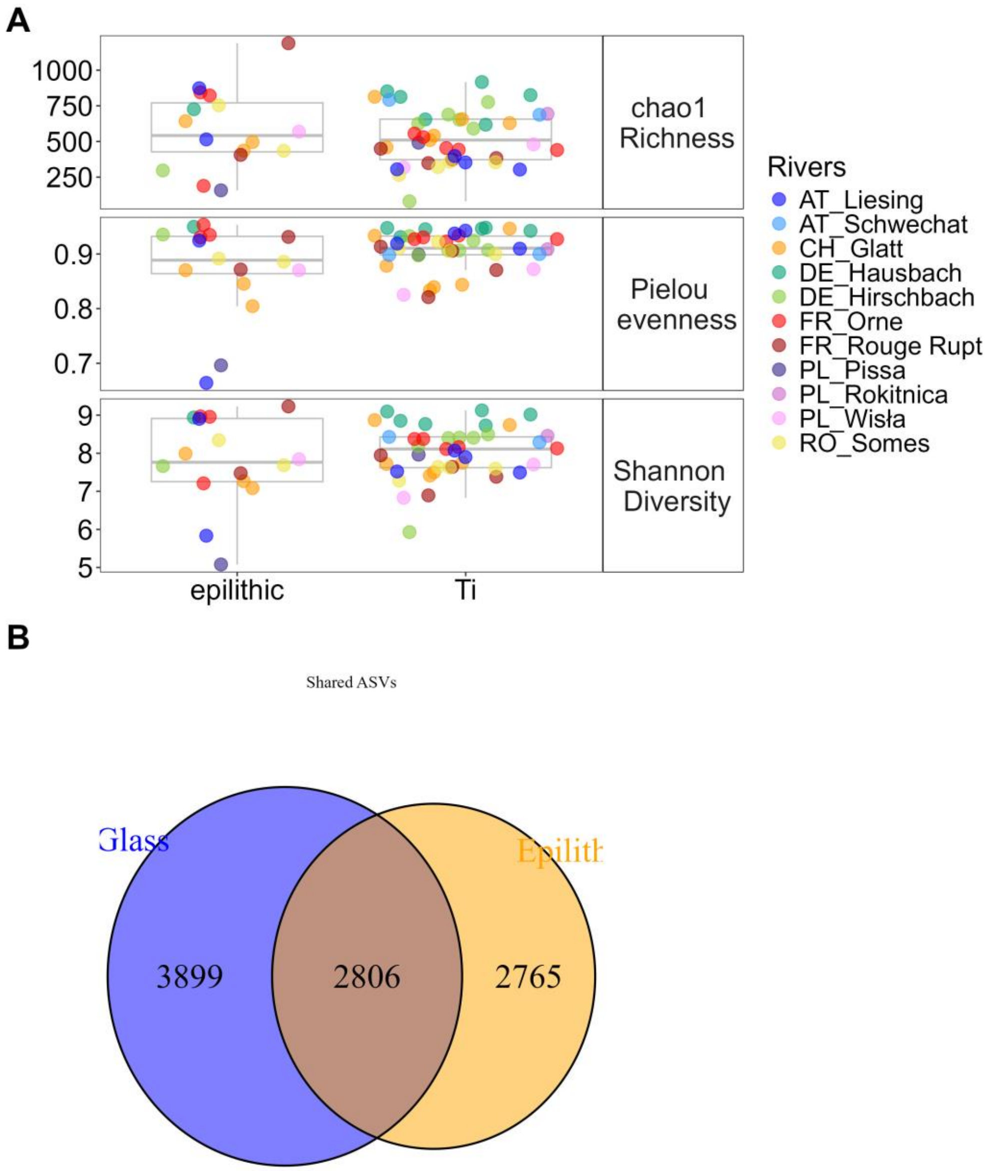


**Supplementary Figure 4.** Comparison of diversity of epilithic biofilms and biofilms established on glass slides (A) Alpha diversity indexes for epilithic biofilm (from Klümper et al [4]) compared to biofilms established on glass slides in this study. (B) Shared and unique ASV numbers and abundance of bacterial taxonomic communities shared and unique to glass or epilithic biofilms from all river sites with paired sampling sites between the two studies.

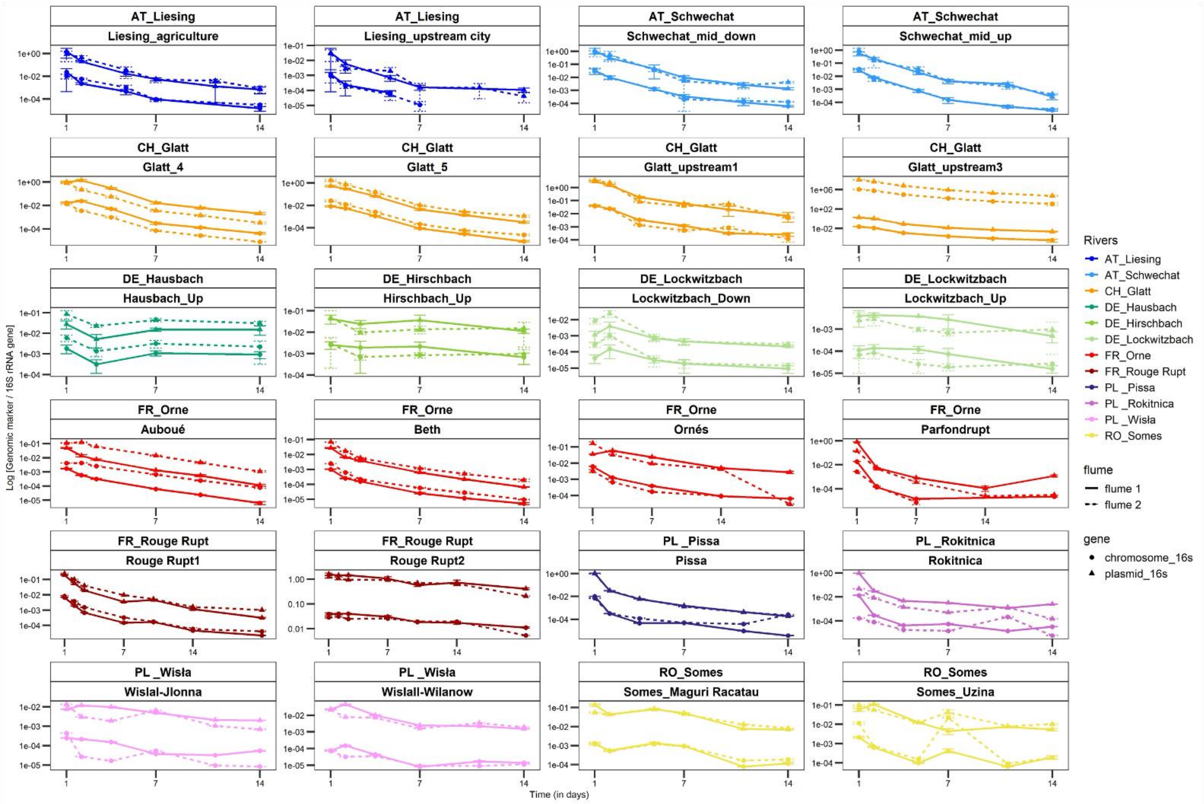


**Supplementary Figure 5.** Invasion dynamics of *E. coli* hosting the pG527 plasmid in all tested flume biofilms. Invader's loss dynamics of the decay kinetics of chromosomal (circular markers) and plasmid (triangular markers) genetic markers in *Escherichia coli*. Persistence of genomic markers normalized to the 16S rRNA gene over a 14-day period of flume incubation. Each point represents the average quantification of three qPCR technical replicates with bars representing the standard deviations. Loss rate per day was calculated using exponential decay kinetics. The full dataset of all flumes and locations is given in Supplementary Table 6.

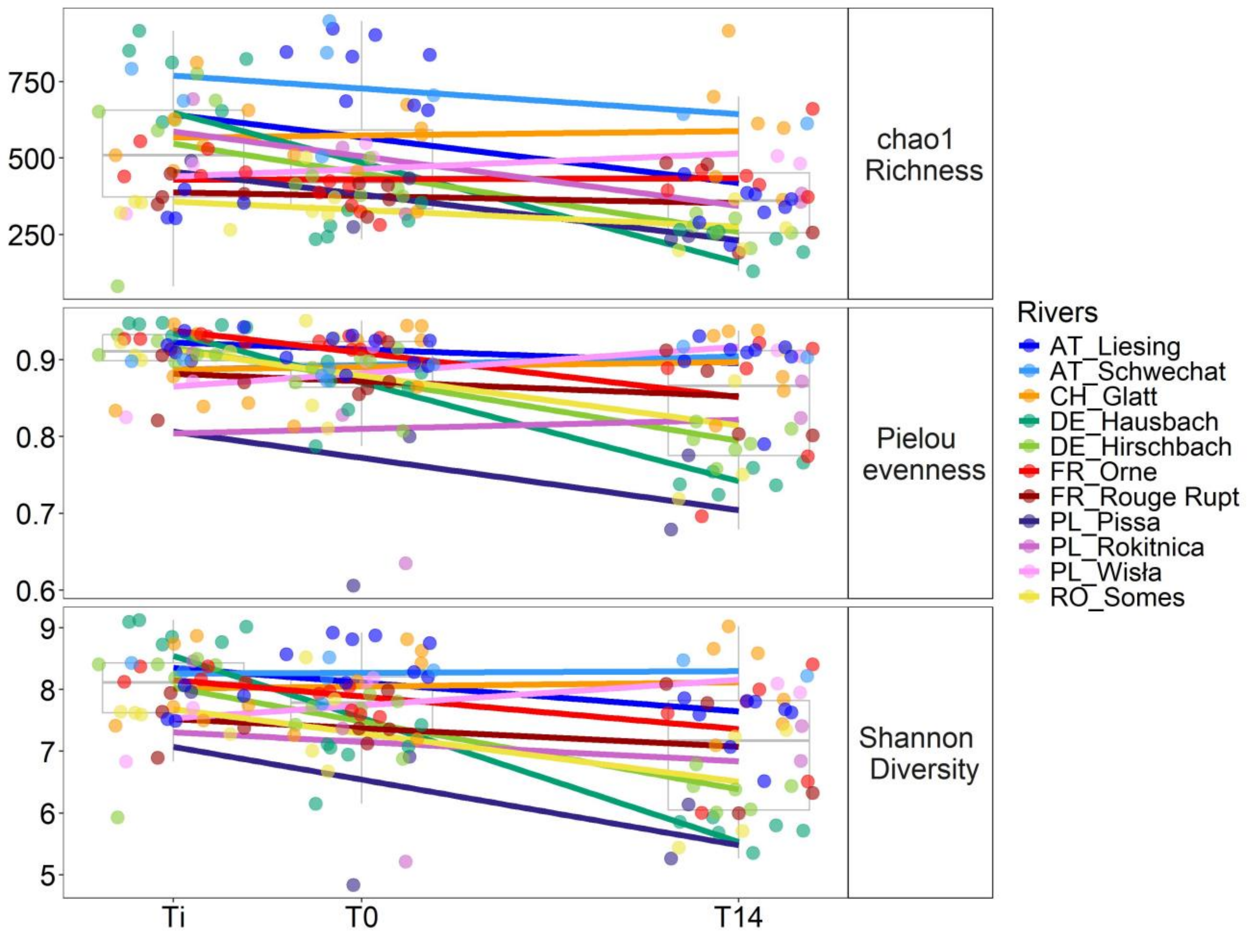


**Supplementary Figure 6.** Alpha diversity indices for biofilms sampled from the river (Ti), after one week of acclimatation in laboratory flumes (T0) and after 14 days of exposure to the model invader, *E. coli*.

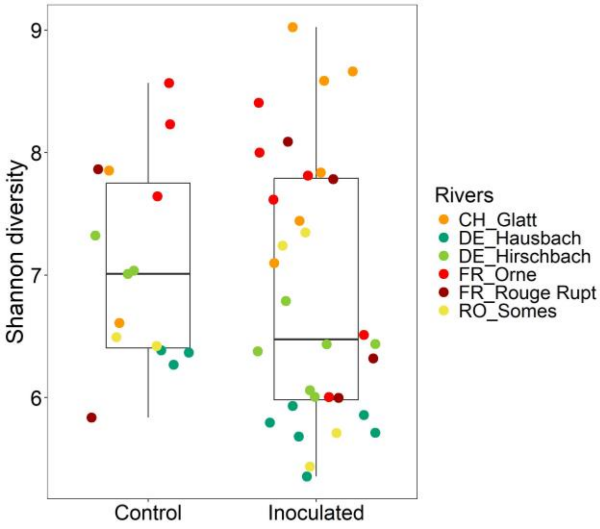


**Supplementary Figure 7.** Alpha Shannon diversity to detect the effect of incubation on the biofilms maintained in control flumes (only part of the river sites were tested) in comparison to those where *E. coli* was added. Colors reflected the country and river where the biofilms were sampled.

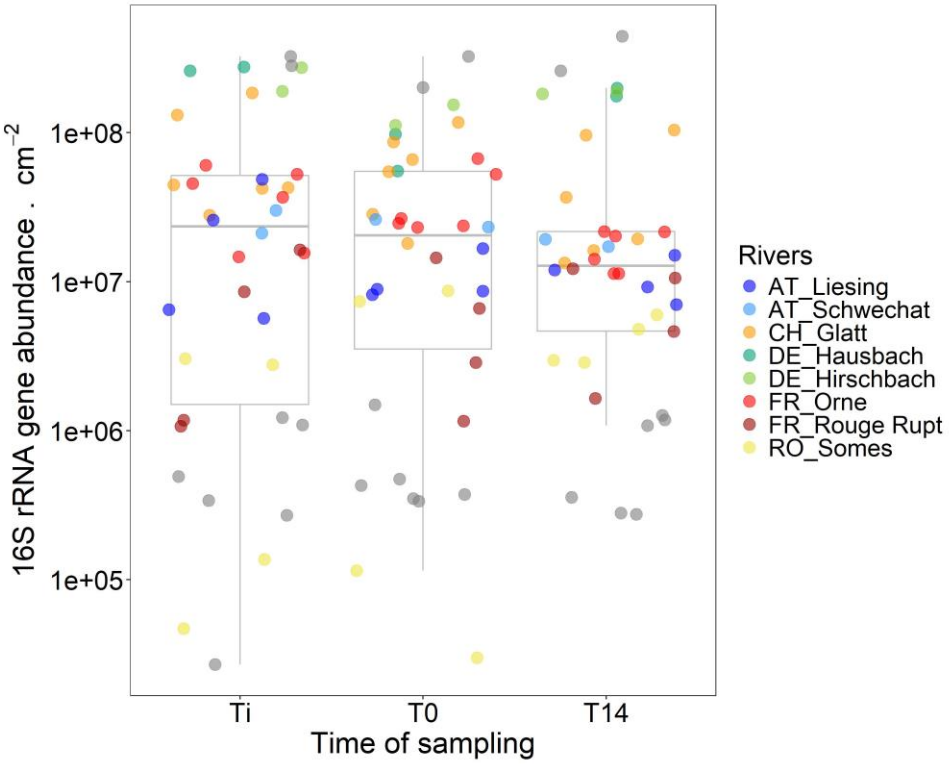


**Supplementary Figure 8.** Quantification of 16S rRNA gene abundance in scraped biofilms from glass slides directly sampled from the river (Ti), after one week of acclimatation in laboratory flumes (T0) and after 14 days of exposure to the model invader, *E. coli*. Colors reflected the country and river where the biofilms were sampled.

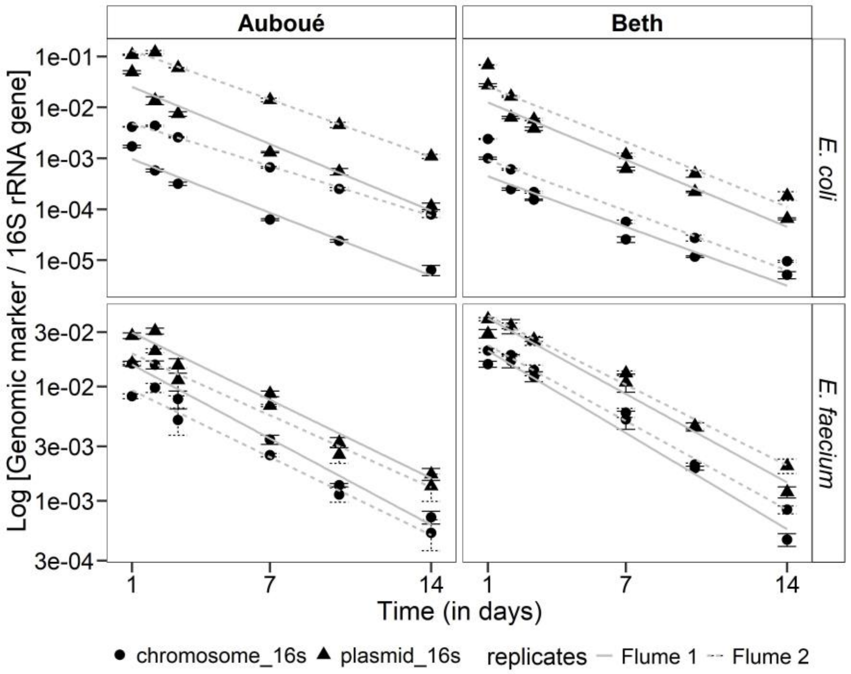


**Supplementary Figure 9.** Invasion dynamics of E. coli hosting the pG527 plasmid and E. faecium hosting the pSF3 plasmid in flume biofilms. Invader's loss dynamics in two river sites over time, showing the same trend of decay kinetics of chromosomal (circular markers) and plasmid (triangular markers) genetic markers in Escherichia coli (upper panel) and Enterococcus faecium (lower panel). Persistence of genomic markers normalized to the 16S rRNA gene over a 14-day period of flume incubation. Each point represents the average quantification of three qPCR technical replicates with bars representing the standard deviations.

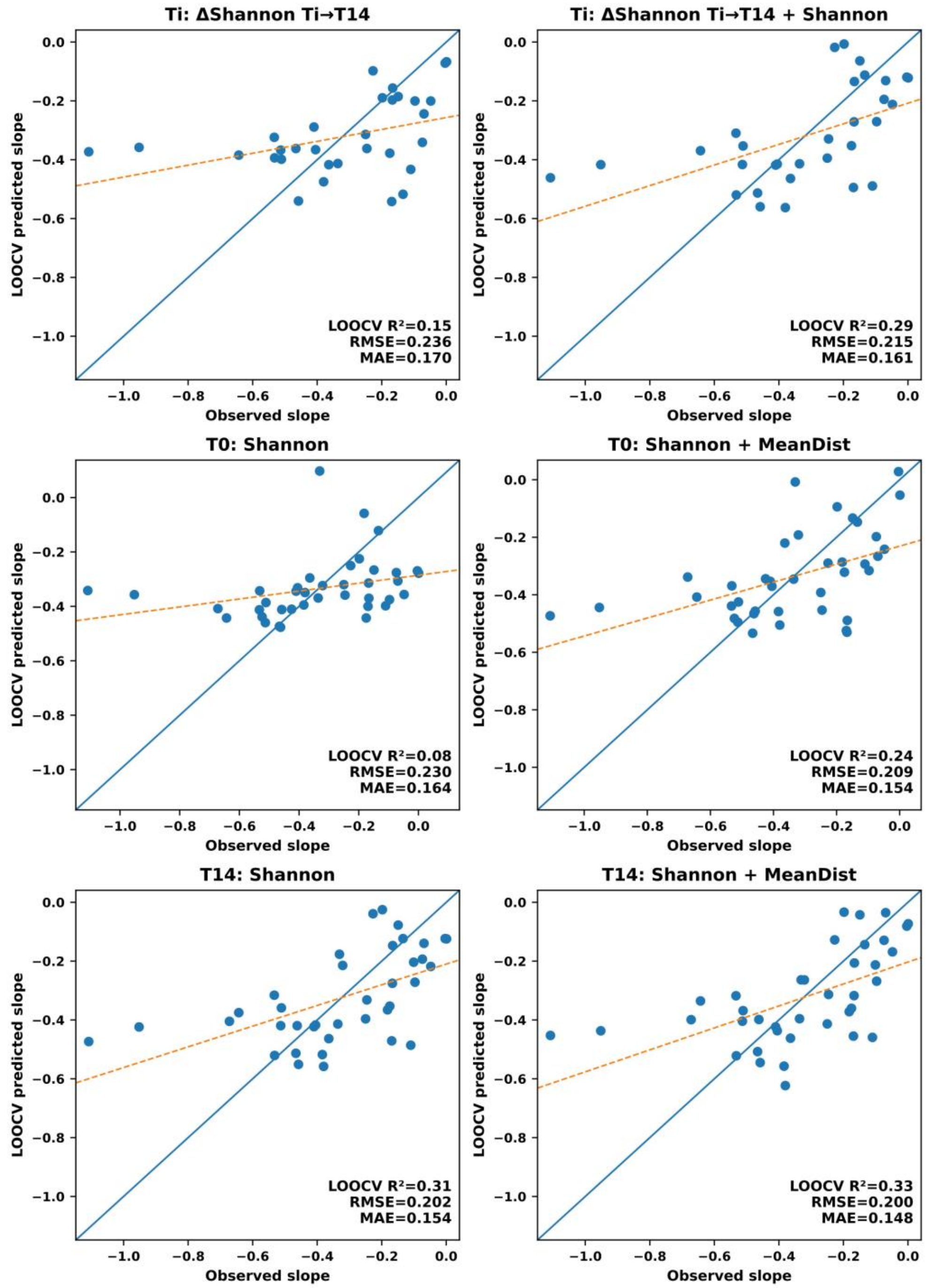


**Supplementary Figure 10.** Validation of top-performing ecological prediction models for ARB invasion resistance. Observed versus predicted invader loss rates for the best-performing ecological prediction models compared to the best single predictor model. Predictions were generated using leave-one-out cross-validation (LOOCV). Solid lines indicate linear fits between observed and predicted values.

### Supplementary Tables

**Supplementary Table 1.** Samples identification and their information regarding their origins, country, rivers, river sites and the respective coordinates.

| Sample names | Country | River | LATITUDE | LONGITUDE |
|---|---|---|---|---|
| PL_Ro_1 | Poland | Rokitnica | 52.0781347 | 20.6966351 |
| PL_Ro_1 | Poland | Rokitnica | 52.148938 | 20.621845 |
| PL_Pi_2 | Poland | Pissa | 53.16117 | 19.53874 |
| PL_Pi_2 | Poland | Pissa | 53.17366 | 19.54804 |
| PL_Wi_3 | Poland | Wisła | 52.170556 | 21.136389 |
| PL_Wi_3 | Poland | Wisła | 52.170556 | 21.136389 |
| DE_Ha_1 | Germany | Hausbach | 50.9141406 | 13.7870833 |
| DE_Ha_1 | Germany | Hausbach | 50.9141406 | 13.7870833 |
| DE_Ha_1 | Germany | Hausbach | 50.9141406 | 13.7870833 |
| DE_Ha_1 | Germany | Hausbach | 50.9141406 | 13.7870833 |
| DE_Ha_1 | Germany | Hausbach | 50.9141406 | 13.7870833 |
| DE_Ha_1 | Germany | Hausbach | 50.9141406 | 13.7870833 |
| DE_Hi_2 | Germany | Hirschbach | 50.9048749 | 13.7518679 |
| DE_Hi_2 | Germany | Hirschbach | 50.9048749 | 13.7518679 |
| DE_Hi_2 | Germany | Hirschbach | 50.9048749 | 13.7518679 |
| DE_Hi_2 | Germany | Hirschbach | 50.9048749 | 13.7518679 |
| DE_Hi_2 | Germany | Hirschbach | 50.9048749 | 13.7518679 |
| DE_Hi_2 | Germany | Hirschbach | 50.9048749 | 13.7518679 |
| DE_Lo_3 | Germany | Lockwitzbach | 50.9605245 | 13.7770674 |
| DE_Lo_3 | Germany | Lockwitzbach | 50.9605245 | 13.7770674 |
| CH_Gl_1 | Switzerland | Glatt | 47.40026 | 9.22152 |
| CH_Gl_1 | Switzerland | Glatt | 47.40026 | 9.22152 |
| CH_Gl_2 | Switzerland | Glatt | 47.39815 | 9.23268 |
| CH_Gl_2 | Switzerland | Glatt | 47.39815 | 9.23268 |
| CH_Gl_3 | Switzerland | Glatt | 47.41404 | 9.20399 |
| CH_Gl_3 | Switzerland | Glatt | 47.41404 | 9.20399 |
| FR_Or_1 | France | Orne | 49.254041 | 5.471382 |
| FR_Or_1 | France | Orne | 49.254041 | 5.471382 |
| FR_Or_2 | France | Orne | 49.160919 | 5.729048 |
| FR_Or_2 | France | Orne | 49.160919 | 5.729048 |
| FR_Or_3 | France | Orne | 49.211258 | 5.976874 |
| FR_Or_3 | France | Orne | 49.211258 | 5.976874 |
| FR_Ro_1 | France | Rouge Rupt | 47.965312 | 6.84458 |
| FR_Ro_1 | France | Rouge Rupt | 47.965312 | 6.84458 |
| FR_Ro_2 | France | Rouge Rupt | 47.974679 | 6.8747736 |
| FR_Ro_2 | France | Rouge Rupt | 47.974679 | 6.8747736 |
| RO_So_1 | Romania | Somes | 46.613407 | 23.124938 |
| RO_So_1 | Romania | Somes | 46.613407 | 23.124938 |
| RO_So_2 | Romania | Somes | 46.686266 | 23.289098 |
| RO_So_2 | Romania | Somes | 46.686266 | 23.289098 |
| AU_Li_1 | Austria | Liesing | 48.137378 | 16.209081 |

| | | | | |
|---|---|---|---|---|
| AU_Li_1 | Austria | Liesing | 48.137378 | 16.209081 |
| AU_Li_1 | Austria | Liesing | 48.137378 | 16.209081 |
| AU_Li_1 | Austria | Liesing | 48.137378 | 16.209081 |
| AU_Li_2 | Austria | Liesing | 48.134789 | 16.172442 |
| AU_Li_2 | Austria | Liesing | 48.134789 | 16.172442 |
| AU_Li_2 | Austria | Liesing | 48.134789 | 16.172442 |
| AU_Li_2 | Austria | Liesing | 48.134789 | 16.172442 |
| AU_Sc_3 | Austria | Schwechat | 48.043642 | 16.092503 |
| AU_Sc_3 | Austria | Schwechat | 48.043642 | 16.092503 |
| AU_Sc_3 | Austria | Schwechat | 48.043642 | 16.092503 |
| AU_Sc_3 | Austria | Schwechat | 48.043642 | 16.092503 |

**Supplementary Table 2: List of samples and their original identifier and NCBI accession ID**

SUBMISSION ONGOING AND LIST TO DO BE ADDED

**Supplementary Table 3: Primers and probes used in this study**

| Strain/plasmid target | Amplicon target | Primer | Sequence | Reference |
|---|---|---|---|---|
| All bacteria | 16S rRNA gene | **338F** | CCTACGGGAGGCAGCAG | [5] |
| | | **518R** | ATTACCGCGGCTGCTGG | |
| *E. coli* CM2372 | Δ*lacZY* locus of EDCM367 | **EDCM-F** | GACAGGTTTCCCGACTGG | [6] |
| | | **EDCM-R** | CGACTTCATTCACCTGACGA | |
| | | **EDCM-P** | FAM-CCGCTTGGAACGGGCTCACT-BHQ1 | |
| pG527 | *nptII* (= *aph(3')II*)) | **pG527-1b** | ATGCAAGAACGGAAATGGAC | [7] |
| | | **pG527-2b/1c** | TTCCGTTCAGGACGCTACTT | |
| | | ***pG527-Pb*** | TCATTTGCCTTTATTAAGTCGAGC | |
| *E. faecium* CM2451 | Δ*sodA* mutation | **EfCom12sodA-1** | TGCAGCAATCGAAAAATATCC | This work |
| | | **EfCom12sodA-2** | TTAGCATGTCCACCCATTCA | |
| | | **EfCom12sodA-P** | TGATATGGATGCTGTTCCAACA | |
| pSF3 | Δ*vanZ* | **pSF3-1** | CTGCTACTGGGAGCTCGTTT | This work |
| | | **pSF3-2** | TTTTCCCCTCACTTCACACC | |
| | | **pSF3-P** | AGATGGAAAACGGGATGTGG | |

**Supplementary Table 4.** Alpha diversity for all flumes and epilithic samples

| samplename | country | country_river | Time | Samples | chao1 Richness | Shannon diversity | Pielou evenness |
|---|---|---|---|---|---|---|---|
| PL_Ro_1 | Poland | PL_Rokitnica | Ti | PL_Ti_Lowdiversityflume1 | 693.850467 | 8.45740119 | 0.90880562 |
| PL_Pi_2 | Poland | PL_Pissa | Ti | PL_Ti_Highdiversityflume2 | 491.652174 | 7.96358716 | 0.89871449 |
| PL_Wi_3 | Poland | PL_Wisła | Ti | PL_Ti_BeforeWWTPflume1 | 318.410256 | 6.83055198 | 0.8253322 |
| PL_Wi_3 | Poland | PL_Wisła | Ti | PL_Ti_BeforeWWTPflumeII | 479.741936 | 7.70270451 | 0.87204805 |
| DE_Ha_1 | Germany | DE_Hausbach | Ti | DE_Ti_LowdiversityFlume1A | 654.882979 | 8.76702734 | 0.94535482 |
| DE_Ha_1 | Germany | DE_Hausbach | Ti | DE_Ti_LowdiversityFlume1B | 812.945946 | 8.85297747 | 0.93093448 |
| DE_Ha_1 | Germany | DE_Hausbach | Ti | DE_Ti_LowdiversityFlume1C | 916.468354 | 9.12590791 | 0.94646951 |
| DE_Ha_1 | Germany | DE_Hausbach | Ti | DE_Ti_LowdiversityFlume2A | 618.013699 | 8.7292323 | 0.94810674 |
| DE_Ha_1 | Germany | DE_Hausbach | Ti | DE_Ti_LowdiversityFlume2B | 851.543478 | 9.09569813 | 0.9476608 |
| DE_Ha_1 | Germany | DE_Hausbach | Ti | DE_Ti_LowdiversityFlume2C | 825.070866 | 9.01630428 | 0.94234382 |
| DE_Hi_2 | Germany | DE_Hirschbach | Ti | DE_Ti_HighdiversityFlume1A | 776.40708 | 8.49556047 | 0.90724459 |
| DE_Hi_2 | Germany | DE_Hirschbach | Ti | DE_Ti_HighdiversityFlume1B | 590.347826 | 8.40607556 | 0.92448038 |
| DE_Hi_2 | Germany | DE_Hirschbach | Ti | DE_Ti_HighdiversityFlume1C | 652.62931 | 8.40874071 | 0.90649288 |
| DE_Hi_2 | Germany | DE_Hirschbach | Ti | DE_Ti_HighdiversityFlume2A | 624.740741 | 8.19046348 | 0.89741741 |
| DE_Hi_2 | Germany | DE_Hirschbach | Ti | DE_Ti_HighdiversityFlume2B | 81 | 5.93008134 | 0.93276175 |
| DE_Hi_2 | Germany | DE_Hirschbach | Ti | DE_Ti_HighdiversityFlume2C | 688.330189 | 8.3983095 | 0.90650996 |
| CH_Gl_1 | Switzerland | CH_Glatt | Ti | CH_Ti_Lowdiversityflume_1 | 510 | 7.41510468 | 0.83364324 |
| CH_Gl_1 | Switzerland | CH_Glatt | Ti | CH_Ti_Lowdiversityflume_2 | 458.862069 | 7.71958547 | 0.87843321 |
| CH_Gl_2 | Switzerland | CH_Glatt | Ti | CH_Ti_Highdiversityflume_1 | 540.72093 | 7.49902783 | 0.83940984 |
| CH_Gl_2 | Switzerland | CH_Glatt | Ti | CH_Ti_Highdiversityflume_2 | 628.774194 | 8.74127871 | 0.94643409 |
| CH_Gl_3 | Switzerland | CH_Glatt | Ti | CH_Ti_BeforeWWTPflume_1 | 656.579365 | 7.74790705 | 0.84377851 |
| CH_Gl_3 | Switzerland | CH_Glatt | Ti | CH_Ti_BeforeWWTPflume_2 | 813.446154 | 8.87154055 | 0.93347046 |
| FR_Or_1 | France | FR_Orne | Ti | FR_J_Ti | 530.727273 | 8.37607971 | 0.93038452 |
| FR_Or_1 | France | FR_Orne | Ti | FR_K_Ti | 442.7 | 8.16239051 | 0.93375523 |
| FR_Or_2 | France | FR_Orne | Ti | FR_H_Ti | 454.153846 | 8.11161727 | 0.92270128 |
| FR_Or_2 | France | FR_Orne | Ti | FR_I_Ti | 555.513889 | 8.37551988 | 0.92717012 |
| FR_Or_3 | France | FR_Orne | Ti | FR_PS_Ti_repeat | 440.258621 | 8.12753571 | 0.9272852 |
| FR_Ro_1 | France | FR_Rouge Rupt | Ti | FR_C_Ti | 372.682927 | 7.6414185 | 0.90595795 |

| | | | | | | | |
|---|---|---|---|---|---|---|---|
| FR_Ro_1 | France | FR_Rouge Rupt | Ti | FR_D_Ti | 449.654546 | 7.94699532 | 0.91340157 |
| FR_Ro_2 | France | FR_Rouge Rupt | Ti | FR_A_Ti | 383.54902 | 7.38226302 | 0.87057244 |
| FR_Ro_2 | France | FR_Rouge Rupt | Ti | FR_B_Ti | 348.508197 | 6.89382177 | 0.82102492 |
| RO_So_1 | Romania | RO_Somes | Ti | RO_Ti_Lowdiversityflume_R1 | 265.4 | 7.28034197 | 0.91068553 |
| RO_So_1 | Romania | RO_Somes | Ti | RO_Ti_Lowdiversityflume_R2 | 354.657895 | 7.59619975 | 0.89970989 |
| RO_So_2 | Romania | RO_Somes | Ti | RO_Ti_High_diversityflume_R1 | 358.037037 | 7.62295697 | 0.90602497 |
| RO_So_2 | Romania | RO_Somes | Ti | RO_Ti_High_diversityflume_R2 | 320.884615 | 7.63887703 | 0.92248382 |
| AU_Li_1 | Austria | AT_Liesing | Ti | AU_Ti_43b_F3A2S4 | 303.5 | 7.4949581 | 0.90975842 |
| AU_Li_1 | Austria | AT_Liesing | Ti | AU_Ti_43b_F4A3S7 | 305.214286 | 7.5223649 | 0.91905675 |
| AU_Li_2 | Austria | AT_Liesing | Ti | AU_Ti_42_F3A2S4 | 397.941177 | 8.06934217 | 0.93749651 |
| AU_Li_2 | Austria | AT_Liesing | Ti | AU_Ti_42_F4A2S4 | 353.461539 | 7.89924508 | 0.94270061 |
| AU_Sc_3 | Austria | AT_Schwechat | Ti | AU_FW1_Flume_1_AEU1_Slide_1 | 792.38843 | 8.4299623 | 0.89815533 |
| AU_Sc_3 | Austria | AT_Schwechat | Ti | AU_FW1_Flume_2_AEU2_Slide_4 | 687.142857 | 8.29116864 | 0.90005093 |
| PL_Ro_1 | Poland | PL_Rokitnica | T0 | PL_T0_Lowdiversityflume1 | 535.010753 | 7.36992369 | 0.82828183 |
| PL_Ro_1 | Poland | PL_Rokitnica | T0 | PL_T0_Lowdiversityflume2 | 316.518519 | 5.2119604 | 0.63487301 |
| PL_Pi_2 | Poland | PL_Pissa | T0 | PL_T0_HighdiversityflumeI | 274.5 | 4.8345805 | 0.60604373 |
| PL_Pi_2 | Poland | PL_Pissa | T0 | PL_T0_HighdiversityflumeII | 433.964912 | 6.91222237 | 0.80000306 |
| PL_Wi_3 | Poland | PL_Wisła | T0 | PL_T0_BeforeWWTPflume1 | 503.285714 | 8.18935443 | 0.91668211 |
| PL_Wi_3 | Poland | PL_Wisła | T0 | PL_T0_BeforeWWTPflumeII | 548.433735 | 8.102583 | 0.89805505 |
| DE_Ha_1 | Germany | DE_Hausbach | T0 | DE_T0_LowdiversityFlume1A | 242.111111 | 7.11561676 | 0.89789087 |
| DE_Ha_1 | Germany | DE_Hausbach | T0 | DE_T0_LowdiversityFlume1B | 295.565217 | 7.07395913 | 0.86374978 |
| DE_Ha_1 | Germany | DE_Hausbach | T0 | DE_T0_LowdiversityFlume1C | 355 | 7.42647288 | 0.88311662 |
| DE_Ha_1 | Germany | DE_Hausbach | T0 | DE_T0_LowdiversityFlume2A | 330.783784 | 6.94752118 | 0.83529837 |
| DE_Ha_1 | Germany | DE_Hausbach | T0 | DE_T0_LowdiversityFlume2B | 277.888889 | 7.0588059 | 0.87280845 |
| DE_Ha_1 | Germany | DE_Hausbach | T0 | DE_T0_LowdiversityFlume2C | 234.636364 | 6.14973519 | 0.78768485 |
| DE_Hi_2 | Germany | DE_Hirschbach | T0 | DE_T0_HighdiversityFlume1A | 500.458333 | 7.9213272 | 0.89841691 |
| DE_Hi_2 | Germany | DE_Hirschbach | T0 | DE_T0_HighdiversityFlume1B | 374.442623 | 6.87784203 | 0.80729292 |
| DE_Hi_2 | Germany | DE_Hirschbach | T0 | DE_T0_HighdiversityFlume1C | 414.466667 | 7.4303258 | 0.87054318 |
| DE_Hi_2 | Germany | DE_Hirschbach | T0 | DE_T0_HighdiversityFlume2A | 400.375 | 7.80766419 | 0.91433645 |
| DE_Hi_2 | Germany | DE_Hirschbach | T0 | DE_T0_HighdiversityFlume2B | 378.478261 | 7.69320635 | 0.90175388 |
| DE_Hi_2 | Germany | DE_Hirschbach | T0 | DE_T0_HighdiversityFlume2C | 442.557692 | 7.71655643 | 0.88941847 |
| CH_Gl_1 | Switzerland | CH_Glatt | T0 | CH_T0_Lowdiversityflume_1 | 457.139535 | 8.13659082 | 0.92519969 |

| | | | | | | | |
|---|---|---|---|---|---|---|---|
| CH_Gl_1 | Switzerland | CH_Glatt | T0 | CH_T0_Lowdiversityflume_2 | 597 | 8.42744503 | 0.92522615 |
| CH_Gl_2 | Switzerland | CH_Glatt | T0 | CH_T0_Highdiversityflume_1 | 510.308824 | 7.26142643 | 0.81335204 |
| CH_Gl_2 | Switzerland | CH_Glatt | T0 | CH_T0_Highdiversityflume_2 | 324.774194 | 7.19627473 | 0.86473647 |
| CH_Gl_3 | Switzerland | CH_Glatt | T0 | CH_T0_BeforeWWTPflume_1 | 575.924051 | 8.62691321 | 0.94390992 |
| CH_Gl_3 | Switzerland | CH_Glatt | T0 | CH_T0_BeforeWWTPflume_2 | 674.625 | 8.81146703 | 0.9445572 |
| FR_Or_1 | France | FR_Orne | T0 | FR_J_T0 | 387.428571 | 7.93206225 | 0.92394839 |
| FR_Or_1 | France | FR_Orne | T0 | FR_K_T0 | 408.117647 | 8.04459521 | 0.93106101 |
| FR_Or_2 | France | FR_Orne | T0 | FR_H_T0 | 425.025 | 7.96815939 | 0.91841849 |
| FR_Or_2 | France | FR_Orne | T0 | FR_I_T0 | 281.545455 | 7.55311691 | 0.92853713 |
| FR_Or_3 | France | FR_Orne | T0 | FR_P_T0_repeat | 345.16129 | 7.66261827 | 0.91305153 |
| FR_Or_3 | France | FR_Orne | T0 | FR_S_T0 | 325.076923 | 7.59364949 | 0.91347966 |
| FR_Ro_1 | France | FR_Rouge Rupt | T0 | FR_C_T0 | 411.977273 | 7.98315741 | 0.92317962 |
| FR_Ro_1 | France | FR_Rouge Rupt | T0 | FR_D_T0 | 364.588235 | 7.35695069 | 0.87137273 |
| FR_Ro_2 | France | FR_Rouge Rupt | T0 | FR_A_T0 | 417.015385 | 7.3706331 | 0.85522203 |
| FR_Ro_2 | France | FR_Rouge Rupt | T0 | FR_B_T0 | 308.241379 | 7.12465362 | 0.86282172 |
| RO_So_1 | Romania | RO_Somes | T0 | RO_T0_Lowdiversityflume_R1 | 314.325 | 6.67651415 | 0.81041347 |
| RO_So_1 | Romania | RO_Somes | T0 | RO_T0_Lowdiversityflume_R2 | 327.410256 | 7.00825847 | 0.84078348 |
| RO_So_2 | Romania | RO_Somes | T0 | RO_T0_High_diversityflume_R1 | 503.06 | 8.51846473 | 0.9510291 |
| RO_So_2 | Romania | RO_Somes | T0 | RO_T0_High_diversityflume_R2 | 372.117647 | 7.84296672 | 0.92802873 |
| AU_Li_1 | Austria | AT_Liesing | T0 | AU_T0_43b_F3A2S5 | 847.891892 | 8.57399035 | 0.90272943 |
| AU_Li_1 | Austria | AT_Liesing | T0 | AU_T0_43b_F3A3S8 | 686.676471 | 8.10802496 | 0.87970582 |
| AU_Li_1 | Austria | AT_Liesing | T0 | AU_T0_43b_F4A2S5 | 672.552632 | 8.28403264 | 0.89600188 |
| AU_Li_1 | Austria | AT_Liesing | T0 | AU_T0_43b_F4A3S8 | 657.185567 | 8.20683204 | 0.89184108 |
| AU_Li_2 | Austria | AT_Liesing | T0 | AU_T0_42_F3A1S2 | 832.53719 | 8.81155008 | 0.93191382 |
| AU_Li_2 | Austria | AT_Liesing | T0 | AU_T0_42_F3A3S8 | 837.75 | 8.75338665 | 0.924761 |
| AU_Li_2 | Austria | AT_Liesing | T0 | AU_T0_42_F4A1S2 | 902.781457 | 8.87455111 | 0.92497924 |
| AU_Li_2 | Austria | AT_Liesing | T0 | AU_T0_42_F4A2S5 | 923.533784 | 8.92190606 | 0.9277659 |
| AU_Sc_3 | Austria | AT_Schwechat | T0 | AU_FW1_F1A1S2_T0 | 504.559322 | 7.76853257 | 0.87981614 |
| AU_Sc_3 | Austria | AT_Schwechat | T0 | AU_FW1_F1A3S8_T0 | 706 | 8.30901297 | 0.89395933 |
| AU_Sc_3 | Austria | AT_Schwechat | T0 | AU_FW1_F2A2S5_T0 | 948.493243 | 8.52052129 | 0.8885989 |
| AU_Sc_3 | Austria | AT_Schwechat | T0 | AU_FW1_F2A3S8_T0 | 845.631579 | 8.1861267 | 0.87298101 |
| PL_Ro_1 | Poland | PL_Rokitnica | T14 | PL_T14_Lowdiversityflumel | 356.765957 | 6.8462908 | 0.82402641 |

| | | | | | | | |
|---|---|---|---|---|---|---|---|
| PL_Ro_1 | Poland | PL_Rokitnica | T14 | PL_T14_LowdiversityflumeII | 383.9375 | 7.40783898 | 0.87193566 |
| PL_Pi_2 | Poland | PL_Pissa | T14 | PL_T14_HighdiversityflumeI | 245.216216 | 6.14140267 | 0.77554094 |
| PL_Pi_2 | Poland | PL_Pissa | T14 | PL_T14_Highdiversityflume2 | 234.918367 | 5.26245039 | 0.67918423 |
| PL_Wi_3 | Poland | PL_Wisła | T14 | PL_T14_BeforeWWTPflumeI | 482.645833 | 7.95149517 | 0.90515944 |
| PL_Wi_3 | Poland | PL_Wisła | T14 | PL_T14_BeforeWWTPflumeII | 507.6 | 8.09705751 | 0.91218859 |
| DE_Ha_1 | Germany | DE_Hausbach | T14 | DE_T14_LowdiversityFlume1A | 265.529412 | 5.85813982 | 0.73811353 |
| DE_Ha_1 | Germany | DE_Hausbach | T14 | DE_T14_LowdiversityFlume1B | 191.928571 | 5.71405846 | 0.76601794 |
| DE_Ha_1 | Germany | DE_Hausbach | T14 | DE_T14_LowdiversityFlume1C | 130.235294 | 5.35647597 | 0.75921496 |
| DE_Ha_1 | Germany | DE_Hausbach | T14 | DE_T14_LowdiversityFlume2A | 236.181818 | 5.79772843 | 0.73665308 |
| DE_Ha_1 | Germany | DE_Hausbach | T14 | DE_T14_LowdiversityFlume2B | 255.607143 | 5.9328093 | 0.75440855 |
| DE_Ha_1 | Germany | DE_Hausbach | T14 | DE_T14_LowdiversityFlume2C | 259.333333 | 5.68411745 | 0.72450764 |
| DE_Hi_2 | Germany | DE_Hirschbach | T14 | DE_T14_HighdiversityFlume1A | 319.666667 | 6.79042548 | 0.81956498 |
| DE_Hi_2 | Germany | DE_Hirschbach | T14 | DE_T14_HighdiversityFlume1B | 278.12 | 6.43630994 | 0.7963612 |
| DE_Hi_2 | Germany | DE_Hirschbach | T14 | DE_T14_HighdiversityFlume1C | 253.923077 | 6.00754288 | 0.75806751 |
| DE_Hi_2 | Germany | DE_Hirschbach | T14 | DE_T14_HighdiversityFlume2A | 255 | 6.43864793 | 0.81005919 |
| DE_Hi_2 | Germany | DE_Hirschbach | T14 | DE_T14_HighdiversityFlume2B | 204.619048 | 6.06001469 | 0.79057392 |
| DE_Hi_2 | Germany | DE_Hirschbach | T14 | DE_T14_HighdiversityFlume2C | 303.146342 | 6.37919338 | 0.7827474 |
| CH_Gl_1 | Switzerland | CH_Glatt | T14 | CH_T14_Lowdiversityflume_1 | 364 | 7.4443657 | 0.87789606 |
| CH_Gl_1 | Switzerland | CH_Glatt | T14 | CH_T14_Lowdiversityflume_2 | 439.264706 | 7.09939253 | 0.81404791 |
| CH_Gl_2 | Switzerland | CH_Glatt | T14 | CH_T14_Highdiversityflume_1 | 613.075949 | 8.58645173 | 0.93817575 |
| CH_Gl_2 | Switzerland | CH_Glatt | T14 | CH_T14_Highdiversityflume_2 | 598.358491 | 7.83916537 | 0.85965742 |
| CH_Gl_3 | Switzerland | CH_Glatt | T14 | CH_T14_BeforeWWTPflume_1 | 701.653061 | 8.66235717 | 0.93128643 |
| CH_Gl_3 | Switzerland | CH_Glatt | T14 | CH_T14_BeforeWWTPflume_2 | 917.110294 | 9.02379625 | 0.93729052 |
| FR_Or_1 | France | FR_Orne | T14 | FR_JT14 | 660.821053 | 8.4072452 | 0.9143521 |
| FR_Or_1 | France | FR_Orne | T14 | FR_KT14 | 412.122449 | 8.00197784 | 0.92194207 |
| FR_Or_2 | France | FR_Orne | T14 | FR_HT14 | 461 | 6.00441185 | 0.69610648 |
| FR_Or_2 | France | FR_Orne | T14 | FR_IT14 | 393.956522 | 7.61704475 | 0.88921278 |
| FR_Or_3 | France | FR_Orne | T14 | FR_PT14 | 441.818182 | 7.81199677 | 0.88861927 |
| FR_Or_3 | France | FR_Orne | T14 | FR_ST14 | 373.125 | 6.5134108 | 0.77415009 |
| FR_Ro_1 | France | FR_Rouge Rupt | T14 | FR_CT14 | 479.722222 | 7.7834195 | 0.88536859 |
| FR_Ro_1 | France | FR_Rouge Rupt | T14 | FR_DT14 | 484.966667 | 8.09055031 | 0.9124051 |
| FR_Ro_2 | France | FR_Rouge Rupt | T14 | FR_AT14 | 190.5 | 5.99776554 | 0.80317118 |

| | | | | | | | |
|---|---|---|---|---|---|---|---|
| FR_Ro_2 | France | FR_Rouge Rupt | T14 | FR_BT14 | 256.583333 | 6.32238485 | 0.80144386 |
| RO_So_1 | Romania | RO_Somes | T14 | RO_T14_Lowdiversityflume_R1 | 198.058824 | 5.43644093 | 0.71889271 |
| RO_So_1 | Romania | RO_Somes | T14 | RO_T14_Lowdiversityflume_R2 | 201.588235 | 5.7123957 | 0.75090675 |
| RO_So_2 | Romania | RO_Somes | T14 | RO_T14_High_diversityflume_R1 | 271.473684 | 7.34873778 | 0.91167535 |
| RO_So_2 | Romania | RO_Somes | T14 | RO_T14_High_diversityflume_R2 | 366.285714 | 7.24217756 | 0.87215424 |
| AU_Li_1 | Austria | AT_Liesing | T14 | AU_T14_43b_F3A1S8 | 365.676471 | 7.62775378 | 0.90389168 |
| AU_Li_1 | Austria | AT_Liesing | T14 | AU_T14_43b_F3A3S5 | 289.214286 | 7.59453056 | 0.93071695 |
| AU_Li_1 | Austria | AT_Liesing | T14 | AU_T14_43b_F4A1S8 | 381.410256 | 7.80480588 | 0.91276307 |
| AU_Li_1 | Austria | AT_Liesing | T14 | AU_T14_43b_F4A3S5 | 321.972973 | 6.51777019 | 0.79023127 |
| AU_Li_2 | Austria | AT_Liesing | T14 | AU_T14_42_F3A2S2 | 447.568182 | 7.85813529 | 0.89825812 |
| AU_Li_2 | Austria | AT_Liesing | T14 | AU_T14_42_F3A3S5 | 339.107143 | 7.67980555 | 0.91651256 |
| AU_Li_2 | Austria | AT_Liesing | T14 | AU_T14_42_F4A1S8 | 386.121951 | 7.81315933 | 0.90970066 |
| AU_Li_2 | Austria | AT_Liesing | T14 | AU_T14_42_F4A2S2 | 214.857143 | 7.07014854 | 0.91328279 |
| AU_Sc_3 | Austria | AT_Schwechat | T14 | AU_FW1_Flume_1_AEU2_Slide_2 | 644.557692 | 8.47795468 | 0.91744899 |
| AU_Sc_3 | Austria | AT_Schwechat | T14 | AU_FW1_Flume_2_AEU3_Slide_5 | 613.064516 | 8.22076003 | 0.90331448 |
| PL_Pi_2 | Poland | PL_Pissa | epilithic | PL_W_2_2 | 157.894737 | 5.07890732 | 0.69625379 |
| PL_Wi_3 | Poland | PL_Wisła | epilithic | PL_W_14_2 | 569.060606 | 7.84211258 | 0.87026022 |
| DE_Ha_1 | Germany | DE_Hausbach | epilithic | D_W_HauUp | 726.16129 | 8.93908194 | 0.94980306 |
| DE_Hi_2 | Germany | DE_Hirschbach | epilithic | D_W_HIR | 297.125 | 7.66258621 | 0.93562275 |
| CH_Gl_1 | Switzerland | CH_Glatt | epilithic | CH_W_BG3 | 496.285714 | 7.08337688 | 0.80455151 |
| CH_Gl_2 | Switzerland | CH_Glatt | epilithic | CH_W_BG1 | 436.181818 | 7.26836856 | 0.84553375 |
| CH_Gl_3 | Switzerland | CH_Glatt | epilithic | CH_W_BG4 | 642.302521 | 7.98951842 | 0.87032653 |
| FR_Or_1 | France | FR_Orne | epilithic | F_W_O1RS | 822.654676 | 8.95768298 | 0.93510675 |
| FR_Or_2 | France | FR_Orne | epilithic | F_W_O3RS | 189 | 7.20959851 | 0.95336781 |
| FR_Or_3 | France | FR_Orne | epilithic | F_W_O5RS | 844.111732 | 8.96759458 | 0.93057423 |
| FR_Ro_1 | France | FR_Rouge Rupt | epilithic | F_W_R1B | 406.405405 | 7.47528223 | 0.87150618 |
| FR_Ro_2 | France | FR_Rouge Rupt | epilithic | F_W_R2B | 1188.35714 | 9.23013117 | 0.93126487 |
| RO_So_1 | Romania | RO_Somes | epilithic | ROU_W_1F4G | 434.567164 | 7.68539233 | 0.88582646 |
| RO_So_2 | Romania | RO_Somes | epilithic | ROU_W_1G3G | 753.17931 | 8.34252754 | 0.89152885 |
| AU_Li_1 | Austria | AT_Liesing | epilithic | AU_W_FW_4_3 | 514.583333 | 5.8340528 | 0.66412013 |
| AU_Li_2 | Austria | AT_Liesing | epilithic | AU_W_FW_4_2 | 873.518987 | 8.90438161 | 0.92453722 |
| DE_Ha_1 | Germany | DE_Hausbach | control_T0 | DE_T0_LowdiversityControlFlumeA | 306 | 6.5415234 | 0.80166992 |

| | | | | | | | |
|---|---|---|---|---|---|---|---|
| DE_Ha_1 | Germany | DE_Hausbach | control_T0 | DE_T0_LowdiversityControlFlumeB | 387.657895 | 7.51690947 | 0.87869909 |
| DE_Ha_1 | Germany | DE_Hausbach | control_T0 | DE_T0_LowdiversityControlFlumeC | 182.235294 | 5.89013565 | 0.78536756 |
| DE_Hi_2 | Germany | DE_Hirschbach | control_T0 | DE_T0_HighdiversityControlFlumeA | 396.763158 | 7.5454733 | 0.87968941 |
| DE_Hi_2 | Germany | DE_Hirschbach | control_T0 | DE_T0_HighdiversityControlFlumeB | 315.55 | 7.3275161 | 0.89310166 |
| DE_Hi_2 | Germany | DE_Hirschbach | control_T0 | DE_T0_HighdiversityControlFlumeC | 368.5 | 7.79811092 | 0.91573342 |
| CH_Gl_1 | Switzerland | CH_Glatt | control_T0 | CH_T0_Lowdiversityflume_control | 381.588235 | 7.70052993 | 0.89895883 |
| CH_Gl_2 | Switzerland | CH_Glatt | control_T0 | CH_T0_Highdiversityflume_control | 414.416667 | 8.17445694 | 0.94335142 |
| FR_Or_1 | France | FR_Orne | control_T0 | FR_L_T0 | 421.526316 | 7.84399486 | 0.91131565 |
| FR_Or_2 | France | FR_Orne | control_T0 | FR_M_T0_repeat | 318 | 7.76229652 | 0.93376709 |
| FR_Or_3 | France | FR_Orne | control_T0 | FR_O_T0_repeat | 290.071429 | 7.38444054 | 0.91427623 |
| FR_Ro_1 | France | FR_Rouge Rupt | control_T0 | FR_E_T0 | 340.6 | 7.63353555 | 0.91336979 |
| FR_Ro_2 | France | FR_Rouge Rupt | control_T0 | FR_F_T0 | 320.6 | 7.14223159 | 0.86059195 |
| RO_So_1 | Romania | RO_Somes | control_T0 | RO_T0_Lowdiversityflume_C | 226 | 7.03680331 | 0.90056314 |
| RO_So_2 | Romania | RO_Somes | control_T0 | RO_T0_High_diversityflume_C | 327.789474 | 7.86325956 | 0.94285384 |
| AU_Sc_3 | Austria | AT_Schwechat | control_T0 | AU_FW1_F3A1S2_Control_T0 | 831.127273 | 8.69059508 | 0.91160123 |
| AU_Sc_3 | Austria | AT_Schwechat | control_T0 | AU_FW1_F3A2S5_Control_T0 | 844.878049 | 8.58766912 | 0.90823606 |
| DE_Ha_1 | Germany | DE_Hausbach | control_T14 | DE_T14_LowdiversityControlFlumeA | 277.5 | 6.38823025 | 0.79412199 |
| DE_Ha_1 | Germany | DE_Hausbach | control_T14 | DE_T14_LowdiversityControlFlumeB | 286.5 | 6.2691178 | 0.7814548 |
| DE_Ha_1 | Germany | DE_Hausbach | control_T14 | DE_T14_LowdiversityControlFlumeC | 254.75 | 6.36808851 | 0.80912242 |
| DE_Hi_2 | Germany | DE_Hirschbach | control_T14 | DE_T14_HighdiversityControlFlumeA | 303 | 7.01099666 | 0.85973673 |
| DE_Hi_2 | Germany | DE_Hirschbach | control_T14 | DE_T14_HighdiversityControlFlumeB | 377.52381 | 7.03661399 | 0.83261605 |
| DE_Hi_2 | Germany | DE_Hirschbach | control_T14 | DE_T14_HighdiversityControlFlumeC | 350.9 | 7.32369349 | 0.87133495 |
| CH_Gl_1 | Switzerland | CH_Glatt | control_T14 | CH_T14_Lowdiversityflume_control | 658.103774 | 7.85535087 | 0.85900978 |
| CH_Gl_2 | Switzerland | CH_Glatt | control_T14 | CH_T14_Highdiversityflume_control | 308.684211 | 6.6107298 | 0.80670237 |
| FR_Or_1 | France | FR_Orne | control_T14 | FR_LT14 | 569 | 8.23328629 | 0.90977062 |
| FR_Or_2 | France | FR_Orne | control_T14 | FR_MT14 | 606.014085 | 8.56894125 | 0.93862149 |
| FR_Or_3 | France | FR_Orne | control_T14 | FR_OT14 | 376.77551 | 7.64516109 | 0.89209917 |
| FR_Ro_1 | France | FR_Rouge Rupt | control_T14 | FR_ET14 | 214.037037 | 5.83791628 | 0.75476864 |
| FR_Ro_2 | France | FR_Rouge Rupt | control_T14 | FR_FT14 | 518.2 | 7.86508509 | 0.88038476 |
| RO_So_1 | Romania | RO_Somes | control_T14 | RO_T14_Lowdiversityflume_C | 267.333333 | 6.49382687 | 0.80724872 |
| RO_So_2 | Romania | RO_Somes | control_T14 | RO_T14_High_diversityflume_C | 262 | 6.42315036 | 0.80010284 |
| DE_Ha_1 | Germany | DE_Hausbach | control_Ti | DE_Ti_LowdiversityControlFlumeA | 724.356436 | 8.94218239 | 0.95034657 |

| | | | | | | | |
|---|---|---|---|---|---|---|---|
| DE_Ha_1 | Germany | DE_Hausbach | control_Ti | DE_Ti_LowdiversityControlFlumeC | 660.191489 | 8.86493576 | 0.95190027 |
| DE_Hi_2 | Germany | DE_Hirschbach | control_Ti | DE_Ti_HighdiversityControlFlumeA | 627.494506 | 8.39021017 | 0.9154714 |
| DE_Hi_2 | Germany | DE_Hirschbach | control_Ti | DE_Ti_HighdiversityControlFlumeB | 801.342593 | 8.60918978 | 0.91516368 |
| DE_Hi_2 | Germany | DE_Hirschbach | control_Ti | DE_Ti_HighdiversityControlFlumeC | 725.71875 | 8.77032278 | 0.9297942 |
| CH_Gl_1 | Switzerland | CH_Glatt | control_Ti | CH_Ti_Lowdiversityflume_control | 368.5 | 7.87599115 | 0.92747614 |
| CH_Gl_2 | Switzerland | CH_Glatt | control_Ti | CH_Ti_Highdiversityflume_control | 502.016949 | 7.59342242 | 0.85724077 |
| RO_So_1 | Romania | RO_Somes | control_Ti | RO_Ti_Lowdiversityflume_C | 325.555556 | 7.649222 | 0.91866772 |
| RO_So_2 | Romania | RO_Somes | control_Ti | RO_Ti_High_diversityflume_C | 388.193548 | 7.86510422 | 0.91817121 |

**Supplementary Table 5.** Ecological descriptors of invasion

| samplename | MeanDist | closeFrac_0.15 | Shannonclose_0.15 | ΔShannon Ti_T14 | ΔShannon Ti_T0 | ΔShannon T0_T14 | ΔEvenness Ti_T14 | ΔEvenness Ti_T0 | ΔEvenness T0_T14 | ΔRichness Ti_T14 | ΔRichness Ti_T0 | ΔRichness T0_T14 |
|---|---|---|---|---|---|---|---|---|---|---|---|---|
| PL_Ro_1 | 0.20927592 | 0.09790476 | 3.59722157 | -1.2349312 | -0.8316168 | -0.4033143 | -0.0999316 | -0.0938176 | -0.006114 | -361 | -164 | -197 |
| PL_Pi_2 | 0.18505879 | 0.27033278 | 3.44317792 | -1.7318908 | -0.7541744 | -0.9777164 | -0.1961027 | -0.1010136 | -0.095089 | -261 | -74 | -187 |
| PL_Wi_3 | 0.19230691 | 0.13521764 | 2.37958524 | 0.64207499 | 0.73756761 | -0.0954926 | 0.04601044 | 0.04602698 | -1.65E-05 | 160 | 212 | -52 |
| PL_Wi_3 | 0.20449218 | 0.13969638 | 2.75118288 | 0.10868688 | 0.13413183 | -0.025445 | 0.01297956 | -0.0053864 | 0.01836596 | 15 | 96 | -81 |
| DE_Ha_1 | 0.21990259 | 0.05917931 | 3.25542369 | -2.0821617 | -1.2380111 | -0.8441506 | -0.212374 | -0.0466723 | -0.1657018 | -415 | -436 | 21 |
| DE_Ha_1 | 0.21132746 | 0.06385623 | 3.31613625 | -2.0526816 | -1.2670756 | -0.7856059 | -0.2079348 | -0.1106034 | -0.0973315 | -387 | -303 | -84 |
| DE_Hi_2 | 0.21330259 | 0.06930458 | 3.18989908 | -1.2423601 | -0.4357058 | -0.8066543 | -0.0748155 | 0.00367409 | -0.0784896 | -481 | -324 | -157 |
| DE_Hi_2 | 0.21042621 | 0.11060295 | 2.82390996 | -1.2624979 | -0.3271231 | -0.9353748 | -0.0810016 | 0.02172584 | -0.1027274 | -385 | -258 | -127 |
| CH_Gl_1 | 0.2077002 | 0.05507901 | 2.85613052 | 0.03206969 | 0.54362203 | -0.5115523 | 0.05344667 | 0.11300784 | -0.0595612 | -154 | -85 | -69 |
| CH_Gl_1 | 0.21423808 | 0.09211912 | 3.04674872 | -0.4113734 | 0.47280911 | -0.8841825 | -0.062695 | 0.03752416 | -0.1002191 | -15 | 138 | -153 |
| CH_Gl_2 | 0.21065319 | 0.04761366 | 2.82778357 | 0.79976991 | 0.63991587 | 0.15985404 | 0.11207132 | 0.11401952 | -0.0019482 | 57 | -45 | 102 |
| CH_Gl_2 | 0.21186125 | 0.0417509 | 3.17724216 | -0.6402657 | -0.6912263 | 0.05096062 | -0.0896692 | -0.0043144 | -0.0853549 | -44 | -324 | 280 |

| | | | | | | | | | | | | |
|---|---|---|---|---|---|---|---|---|---|---|---|---|
| AU_Li_1 | 0.22037698 | 0.10089378 | 2.77781766 | -0.0156994 | | -0.8467198 | -0.026138 | -0.0311886 | 0.00505059 | 48 | 626 | -578 |
| AU_Li_2 | 0.21771189 | 0.05280112 | 2.74899632 | -0.155173 | 0.56494778 | -0.7201208 | -0.0434179 | -0.0244579 | -0.0189599 | 48 | 458 | -410 |
| AU_Sc_3 | 0.20544006 | 0.07030952 | 2.76905251 | 0.00875767 | -0.4841292 | 0.49288685 | 0.0292203 | -0.0140761 | 0.04329642 | -146 | -286 | 140 |
| FR_Or_1 | 0.20750834 | 0.08962881 | 3.28213948 | 0.06194276 | -0.3146044 | 0.37654718 | -0.0210818 | -0.0020188 | -0.019063 | 129 | -150 | 279 |
| FR_Or_1 | 0.21341503 | 0.07718599 | 3.0565617 | -0.0997695 | -0.0761485 | -0.0236209 | -0.011768 | -0.0025529 | -0.0092151 | -13 | -27 | 14 |
| FR_Or_2 | 0.20285966 | 0.13845798 | 3.39968727 | -1.4641613 | -0.1350785 | -1.3290828 | -0.2472405 | -0.0102576 | -0.2369829 | 37 | -35 | 72 |
| FR_Or_2 | 0.20337908 | 0.13601732 | 3.50066111 | -0.5243947 | -0.6157693 | 0.09137461 | -0.0374914 | 0.00492665 | -0.0424181 | -155 | -281 | 126 |
| FR_Or_3 | 0.22618159 | 0.13943307 | 3.46995383 | -0.1892523 | -0.2955694 | 0.10631713 | -0.0431941 | -0.0102898 | -0.0329043 | 39 | -100 | 139 |
| FR_Ro_1 | 0.2044767 | 0.1056867 | 2.71735689 | 0.06539359 | 0.21396883 | -0.1485752 | -0.0261779 | 0.01888687 | -0.0450648 | 101 | 42 | 59 |
| FR_Ro_1 | 0.20534424 | 0.08887594 | 2.77782244 | 0.09596315 | -0.4398005 | 0.53576364 | 0.00436494 | -0.0359861 | 0.04035101 | 35 | -99 | 134 |
| FR_Ro_2 | 0.21094518 | 0.01136615 | 1.7169957 | -0.9858431 | -0.0204399 | -0.9654032 | -0.0657535 | -0.016666 | -0.0490875 | -196 | 36 | -232 |
| FR_Ro_2 | 0.20191693 | 0.01074807 | 1.59269907 | -0.4011695 | 0.11286382 | -0.5140333 | -0.0187737 | 0.04844271 | -0.0672164 | -110 | -65 | -45 |
| RO_So_1 | 0.21306009 | 0.05506142 | 1.95480676 | -1.2645307 | -0.392389 | -0.8721417 | -0.1940312 | -0.1031275 | -0.0909037 | -59 | 59 | -118 |
| RO_So_1 | 0.21609801 | 0.0575233 | 2.44891376 | -1.3091035 | -0.3637672 | -0.9453363 | -0.1591606 | -0.0623834 | -0.0967772 | -133 | -1 | -132 |
| RO_So_2 | 0.20141569 | 0.15055448 | 2.58040169 | -0.1712248 | 0.67154467 | -0.8427695 | 0.00980398 | 0.05318335 | -0.0433794 | -77 | 159 | -236 |
| RO_So_2 | 0.20349657 | 0.12980621 | 2.72850139 | -0.247019 | 0.19243637 | -0.4394554 | -0.0492504 | 0.00678537 | -0.0560358 | 13 | 56 | -43 |
| PL_Ro_1 | 0.18903192 | 0.05813157 | 3.31858498 | -1.2349312 | -0.8316168 | -0.4033143 | -0.0999316 | -0.0938176 | -0.006114 | -361 | -164 | -197 |
| PL_Ro_1 | 0.17688912 | 0.03847526 | 2.22710749 | | | | | | | | | |
| PL_Pi_2 | 0.18210096 | 0.09457798 | 2.57327385 | | | | | | | | | |
| PL_Pi_2 | 0.1984128 | 0.09243111 | 2.63378459 | -1.7318908 | -0.7541744 | -0.9777164 | -0.1961027 | -0.1010136 | -0.095089 | -261 | -74 | -187 |
| PL_Wi_3 | 0.19992052 | 0.1704252 | 3.47167696 | 0.64207499 | 0.73756761 | -0.0954926 | 0.04601044 | 0.04602698 | -1.65E-05 | 160 | 212 | -52 |
| PL_Wi_3 | 0.21890482 | 0.09028222 | 3.45992522 | 0.10868688 | 0.13413183 | -0.025445 | 0.01297956 | -0.0053864 | 0.01836596 | 15 | 96 | -81 |

| | | | | | | | | | | | | |
|---|---|---|---|---|---|---|---|---|---|---|---|---|
| DE_Ha_1 | 0.22247274 | 0.09949367 | 2.78962798 | -2.0821617 | -1.2380111 | -0.8441506 | -0.212374 | -0.0466723 | -0.1657018 | -415 | -436 | 21 |
| DE_Ha_1 | 0.22699386 | 0.04872495 | 2.36446225 | -2.0526816 | -1.2670756 | -0.7856059 | -0.2079348 | -0.1106034 | -0.0973315 | -387 | -303 | -84 |
| DE_Hi_2 | 0.2136054 | 0.09691161 | 2.77682139 | -1.2423601 | -0.4357058 | -0.8066543 | -0.0748155 | 0.00367409 | -0.0784896 | -481 | -324 | -157 |
| DE_Hi_2 | 0.21774446 | 0.07524893 | 2.12141295 | -1.2624979 | -0.3271231 | -0.9353748 | -0.0810016 | 0.02172584 | -0.1027274 | -385 | -258 | -127 |
| CH_Gl_1 | 0.20620167 | 0.12116228 | 3.1261509 | 0.03206969 | 0.54362203 | -0.5115523 | 0.05344667 | 0.11300784 | -0.0595612 | -154 | -85 | -69 |
| CH_Gl_1 | 0.20885082 | 0.08093164 | 3.09779374 | -0.4113734 | 0.47280911 | -0.8841825 | -0.062695 | 0.03752416 | -0.1002191 | -15 | 138 | -153 |
| CH_Gl_2 | 0.2070245 | 0.12485108 | 3.48578748 | 0.79976991 | 0.63991587 | 0.15985404 | 0.11207132 | 0.11401952 | -0.0019482 | 57 | -45 | 102 |
| CH_Gl_2 | 0.20632055 | 0.11001289 | 2.97076564 | -0.6402657 | -0.6912263 | 0.05096062 | -0.0896692 | -0.0043144 | -0.0853549 | -44 | -324 | 280 |
| AU_Li_1 | 0.21602998 | 0.09138688 | 3.21435332 | -0.0156994 | 0.83102043 | -0.8467198 | -0.026138 | -0.0311886 | 0.00505059 | 48 | 626 | -578 |
| AU_Li_1 | 0.21685895 | 0.07912153 | 3.16384275 | | | | | | | | | |
| AU_Li_2 | 0.21071627 | 0.11163464 | 3.29477329 | -0.155173 | 0.56494778 | -0.7201208 | -0.0434179 | -0.0244579 | -0.0189599 | 48 | 458 | -410 |
| AU_Li_2 | 0.21523705 | 0.07620609 | 3.38192933 | | | | | | | | | |
| AU_Sc_3 | 0.2049535 | 0.07490872 | 2.03840364 | 0.00875767 | -0.4841292 | 0.49288685 | 0.0292203 | -0.0140761 | 0.04329642 | -146 | -286 | 140 |
| AU_Sc_3 | 0.20885173 | 0.08138756 | 2.69985521 | | | | | | | | | |
| FR_Or_1 | 0.19275517 | 0.19392795 | 3.27060038 | 0.06194276 | -0.3146044 | 0.37654718 | -0.0210818 | -0.0020188 | -0.019063 | 129 | -150 | 279 |
| FR_Or_1 | 0.20020466 | 0.153326 | 3.11121724 | -0.0997695 | -0.0761485 | -0.0236209 | -0.011768 | -0.0025529 | -0.0092151 | -13 | -27 | 14 |
| FR_Or_2 | 0.19868915 | 0.14045821 | 3.22823704 | -1.4641613 | -0.1350785 | -1.3290828 | -0.2472405 | -0.0102576 | -0.2369829 | 37 | -35 | 72 |
| FR_Or_2 | 0.19951053 | 0.14381354 | 2.93007 | -0.5243947 | -0.6157693 | 0.09137461 | -0.0374914 | 0.00492665 | -0.0424181 | -155 | -281 | 126 |
| FR_Or_3 | 0.20307309 | 0.17592735 | 3.443292 | -0.1892523 | -0.2955694 | 0.10631713 | -0.0431941 | -0.0102898 | -0.0329043 | 39 | -100 | 139 |
| FR_Or_3 | 0.2168404 | 0.11535836 | 3.12445016 | | | | | | | | | |
| FR_Ro_1 | 0.21038843 | 0.07574305 | 2.68293153 | 0.06539359 | 0.21396883 | -0.1485752 | -0.0261779 | 0.01888687 | -0.0450648 | 101 | 42 | 59 |
| FR_Ro_1 | 0.21075165 | 0.05947164 | 2.60405616 | 0.09596315 | -0.4398005 | 0.53576364 | 0.00436494 | -0.0359861 | 0.04035101 | 35 | -99 | 134 |

| | | | | | | | | | | | | |
|---|---|---|---|---|---|---|---|---|---|---|---|---|
| FR_Ro_2 | 0.20814392 | 0.01883059 | 2.30245021 | -0.9858431 | -0.0204399 | -0.9654032 | -0.0657535 | -0.016666 | -0.0490875 | -196 | 36 | -232 |
| FR_Ro_2 | 0.20958497 | 0.01824561 | 1.92071893 | -0.4011695 | 0.11286382 | -0.5140333 | -0.0187737 | 0.04844271 | -0.0672164 | -110 | -65 | -45 |
| RO_So_1 | 0.21091989 | 0.03345686 | 2.38974002 | -1.2645307 | -0.392389 | -0.8721417 | -0.1940312 | -0.1031275 | -0.0909037 | -59 | 59 | -118 |
| RO_So_1 | 0.21371466 | 0.0275797 | 2.36349804 | -1.3091035 | -0.3637672 | -0.9453363 | -0.1591606 | -0.0623834 | -0.0967772 | -133 | -1 | -132 |
| RO_So_2 | 0.22245682 | 0.0758427 | 2.66890631 | -0.1712248 | 0.67154467 | -0.8427695 | 0.00980398 | 0.05318335 | -0.0433794 | -77 | 159 | -236 |
| RO_So_2 | 0.1998261 | 0.0958273 | 3.16274799 | -0.247019 | 0.19243637 | -0.4394554 | -0.0492504 | 0.00678537 | -0.0560358 | 13 | 56 | -43 |
| PL_Ro_1 | 0.19726652 | 0.12361131 | 2.24443647 | -1.2349312 | -0.8316168 | -0.4033143 | -0.0999316 | -0.0938176 | -0.006114 | -361 | -164 | -197 |
| PL_Ro_1 | 0.20724501 | 0.10572988 | 2.6700673 | | | | | | | | | |
| PL_Pi_2 | 0.17984891 | 0.13199961 | 2.24594892 | | | | | | | | | |
| PL_Pi_2 | 0.17763549 | 0.07162233 | 2.58847462 | -1.7318908 | -0.7541744 | -0.9777164 | -0.1961027 | -0.1010136 | -0.095089 | -261 | -74 | -187 |
| PL_Wi_3 | 0.21503373 | 0.04174625 | 2.76112513 | 0.64207499 | 0.73756761 | -0.0954926 | 0.04601044 | 0.04602698 | -1.65E-05 | 160 | 212 | -52 |
| PL_Wi_3 | 0.21947669 | 0.03329892 | 2.56424266 | 0.10868688 | 0.13413183 | -0.025445 | 0.01297956 | -0.0053864 | 0.01836596 | 15 | 96 | -81 |
| DE_Ha_1 | 0.22077684 | 0.02820465 | 2.00626609 | -2.0821617 | -1.2380111 | -0.8441506 | -0.212374 | -0.0466723 | -0.1657018 | -415 | -436 | 21 |
| DE_Ha_1 | 0.21806241 | 0.00978109 | 1.25319074 | -2.0526816 | -1.2670756 | -0.7856059 | -0.2079348 | -0.1106034 | -0.0973315 | -387 | -303 | -84 |
| DE_Hi_2 | 0.2088749 | 0.06105458 | 2.18837971 | -1.2423601 | -0.4357058 | -0.8066543 | -0.0748155 | 0.00367409 | -0.0784896 | -481 | -324 | -157 |
| DE_Hi_2 | 0.22286056 | 0.01126656 | 1.24426093 | -1.2624979 | -0.3271231 | -0.9353748 | -0.0810016 | 0.02172584 | -0.1027274 | -385 | -258 | -127 |
| CH_Gl_1 | 0.20629792 | 0.06721738 | 2.8012104 | 0.03206969 | 0.54362203 | -0.5115523 | 0.05344667 | 0.11300784 | -0.0595612 | -154 | -85 | -69 |
| CH_Gl_1 | 0.20800427 | 0.0504796 | 2.66299347 | -0.4113734 | 0.47280911 | -0.8841825 | -0.062695 | 0.03752416 | -0.1002191 | -15 | 138 | -153 |
| CH_Gl_2 | 0.21281121 | 0.04517656 | 2.89510183 | 0.79976991 | 0.63991587 | 0.15985404 | 0.11207132 | 0.11401952 | -0.0019482 | 57 | -45 | 102 |
| CH_Gl_2 | 0.20953484 | 0.04435454 | 3.19418918 | -0.6402657 | -0.6912263 | 0.05096062 | -0.0896692 | -0.0043144 | -0.0853549 | -44 | -324 | 280 |
| AU_Li_1 | 0.21868809 | 0.04910317 | 2.48359613 | -0.0156994 | 0.83102043 | -0.8467198 | -0.026138 | -0.0311886 | 0.00505059 | 48 | 626 | -578 |
| AU_Li_1 | 0.21115529 | 0.05492894 | 2.50525084 | | | | | | | | | |

| | | | | | | | | | | | | |
|---|---|---|---|---|---|---|---|---|---|---|---|---|
| AU_Li_2 | 0.21387557 | 0.06679372 | 2.95966063 | -0.155173 | 0.56494778 | -0.7201208 | -0.0434179 | -0.0244579 | -0.0189599 | 48 | 458 | -410 |
| AU_Li_2 | 0.21544049 | 0.07504583 | 2.87893021 | | | | | | | | | |
| AU_Sc_3 | 0.21062996 | 0.05287085 | 2.76242207 | 0.00875767 | -0.4841292 | 0.49288685 | 0.0292203 | -0.0140761 | 0.04329642 | -146 | -286 | 140 |
| FR_Or_1 | 0.21555783 | 0.09028514 | 2.99964833 | 0.06194276 | -0.3146044 | 0.37654718 | -0.0210818 | -0.0020188 | -0.019063 | 129 | -150 | 279 |
| FR_Or_1 | 0.20584889 | 0.05023626 | 2.7519989 | -0.0997695 | -0.0761485 | -0.0236209 | -0.011768 | -0.0025529 | -0.0092151 | -13 | -27 | 14 |
| FR_Or_2 | 0.19104749 | 0.04896137 | 3.26330791 | -1.4641613 | -0.1350785 | -1.3290828 | -0.2472405 | -0.0102576 | -0.2369829 | 37 | -35 | 72 |
| FR_Or_2 | 0.20479315 | 0.0680834 | 3.0052854 | -0.5243947 | -0.6157693 | 0.09137461 | -0.0374914 | 0.00492665 | -0.0424181 | -155 | -281 | 126 |
| FR_Or_3 | 0.20436116 | 0.07640198 | 3.15298651 | -0.1892523 | -0.2955694 | 0.10631713 | -0.0431941 | -0.0102898 | -0.0329043 | 39 | -100 | 139 |
| FR_Or_3 | 0.19282438 | 0.02347664 | 2.79389342 | | | | | | | | | |
| FR_Ro_1 | 0.21473214 | 0.05433326 | 2.46989893 | 0.06539359 | 0.21396883 | -0.1485752 | -0.0261779 | 0.01888687 | -0.0450648 | 101 | 42 | 59 |
| FR_Ro_1 | 0.21056498 | 0.06512014 | 2.49207148 | 0.09596315 | -0.4398005 | 0.53576364 | 0.00436494 | -0.0359861 | 0.04035101 | 35 | -99 | 134 |
| FR_Ro_2 | 0.23287904 | 0.00446278 | 1.08877586 | -0.9858431 | -0.0204399 | -0.9654032 | -0.0657535 | -0.016666 | -0.0490875 | -196 | 36 | -232 |
| FR_Ro_2 | 0.22582236 | 0.00316909 | 1.37741724 | -0.4011695 | 0.11286382 | -0.5140333 | -0.0187737 | 0.04844271 | -0.0672164 | -110 | -65 | -45 |
| RO_So_1 | 0.20305396 | 0.00878202 | 1.41517429 | -1.2645307 | -0.392389 | -0.8721417 | -0.1940312 | -0.1031275 | -0.0909037 | -59 | 59 | -118 |
| RO_So_1 | 0.21434699 | 0.00395452 | 0.52532132 | -1.3091035 | -0.3637672 | -0.9453363 | -0.1591606 | -0.0623834 | -0.0967772 | -133 | -1 | -132 |
| RO_So_2 | 0.20680872 | 0.11051144 | 2.8125445 | -0.1712248 | 0.67154467 | -0.8427695 | 0.00980398 | 0.05318335 | -0.0433794 | -77 | 159 | -236 |
| RO_So_2 | 0.21391705 | 0.07863025 | 2.79978718 | -0.247019 | 0.19243637 | -0.4394554 | -0.0492504 | 0.00678537 | -0.0560358 | 13 | 56 | -43 |

**Supplementary Table 6.** Linear regressions for the decay kinetics of chromosomal marker for *E. coli* in all tested biofilm flumes. The estimated slopes, $R^2$ explaining the qPCR data points over the 14 days of incubation, and the calculated loss rate represented as the log loss per day for the chromosome marker (log_day_chromosome) and the plasmid marker (log_day_plasmid).

| samplename | country_river | slope_plasmid | r2_plasmid | log_day_plasmid | slope_chromosome | r2_chromosome | log_day_chromosome |
|---|---|---|---|---|---|---|---|
| PL_Ro_1 | PL _Rokitnica | -0.197 | 0.633 | 0.08555601 | -0.167 | 0.557 | 0.072527178 |
| PL_Ro_1 | PL _Rokitnica | -0.3 | 0.8698 | 0.13028835 | -0.182 | 0.79 | 0.079041596 |
| PL_Pi_2 | PL _Pissa | -0.407 | 0.9591 | 0.17675785 | -0.331 | 0.9235 | 0.143751474 |
| PL_Pi_2 | PL _Pissa | -0.469 | 0.9502 | 0.20368411 | -0.227 | 0.8729 | 0.098584847 |
| PL_Wi_3 | PL _Wisła | -0.254 | 0.8007 | 0.1103108 | -0.169 | 0.525 | 0.073395767 |
| PL_Wi_3 | PL _Wisła | -0.108 | 0.5503 | 0.0469038 | -0.11 | 0.6119 | 0.047772393 |
| DE_Ha_1 | DE_Hausbach | 0.003 | 0.0006 | -0.0013029 | 0.0009 | 0.00005 | -0.000390865 |
| DE_Ha_1 | DE_Hausbach | -0.041 | 0.0957 | 0.01780607 | -0.0035 | 0.1724 | 0.001520031 |
| DE_Hi_2 | DE_Hirschbach | -0.087 | 0.7153 | 0.03778362 | -0.097 | 0.8454 | 0.042126565 |
| DE_Hi_2 | DE_Hirschbach | -0.047 | 0.1465 | 0.02041184 | -0.048 | 0.1854 | 0.020846135 |
| DE_Lo_3 | DE_Lockwitzbach | -0.167 | 0.8968 | 0.07252718 | -0.162 | 0.8869 | 0.070355706 |
| DE_Lo_3 | DE_Lockwitzbach | -0.085 | 0.4738 | 0.03691503 | -0.083 | 0.4516 | 0.036046442 |
| CH_Gl_1 | CH_Glatt | -0.534 | 0.9109 | 0.23191325 | -0.511 | 0.9208 | 0.22192448 |
| CH_Gl_1 | CH_Glatt | -0.585 | 0.9384 | 0.25406227 | -0.533 | 0.9436 | 0.231478959 |
| CH_Gl_2 | CH_Glatt | -0.406 | 0.8976 | 0.17632356 | -0.458 | 0.9355 | 0.198906873 |
| CH_Gl_2 | CH_Glatt | -0.464 | 0.8331 | 0.20151264 | -0.41 | 0.8222 | 0.178060738 |
| CH_Gl_3 | CH_Glatt | -0.54 | 0.9409 | 0.23451902 | -0.522 | 0.9395 | 0.22670172 |
| CH_Gl_3 | CH_Glatt | -0.598 | 0.9502 | 0.2597081 | -0.572 | 0.9457 | 0.248416444 |
| AU_Li_1 | AT_Liesing | -0.906 | 0.957 | 0.3934708 | -0.643 | 0.8658 | 0.279251352 |
| AU_Li_1 | AT_Liesing | -0.723 | 0.8534 | 0.31399491 | -0.672 | 0.9613 | 0.291845892 |
| AU_Li_2 | AT_Liesing | -0.557 | 0.8996 | 0.24190203 | -0.513 | 0.879 | 0.222793069 |
| AU_Li_2 | AT_Liesing | -0.519 | 0.9152 | 0.22539884 | -0.462 | 0.8811 | 0.200644051 |
| AU_Sc_3 | AT_Schwechat | -0.552 | 0.9469 | 0.23973055 | -0.532 | 0.9228 | 0.231044664 |
| AU_Sc_3 | AT_Schwechat | -0.559 | 0.9404 | 0.24277062 | -0.524 | 0.9046 | 0.227570309 |
| FR_Or_1 | FR_Orne | -1.051 | 0.8201 | 0.4564435 | -1.108 | 0.8475 | 0.481198286 |
| FR_Or_1 | FR_Orne | -0.942 | 0.9309 | 0.4091054 | -0.952 | 0.9733 | 0.413448347 |
| FR_Or_2 | FR_Orne | -0.173 | 0.9597 | 0.07513295 | -0.166 | 0.9169 | 0.072092884 |

| | | | | | | | |
|---|---|---|---|---|---|---|---|
| FR_Or_2 | FR_Orne | -0.257 | 0.8423 | 0.11161368 | -0.25 | 0.8208 | 0.10857362 |
| FR_Or_3 | FR_Orne | -0.431 | 0.9687 | 0.18718092 | -0.405 | 0.9733 | 0.175889265 |
| FR_Or_3 | FR_Orne | -0.37 | 0.9929 | 0.16068896 | -0.321 | 0.9939 | 0.139408529 |
| FR_Ro_1 | FR_Rouge Rupt | -0.344 | 0.8625 | 0.1493973 | -0.336 | 0.8712 | 0.145922946 |
| FR_Ro_1 | FR_Rouge Rupt | -0.36 | 0.9498 | 0.15634601 | -0.364 | 0.9544 | 0.158083191 |
| FR_Ro_2 | FR_Rouge Rupt | -0.068 | 0.8929 | 0.02953203 | -0.069 | 0.9609 | 0.029966319 |
| FR_Ro_2 | FR_Rouge Rupt | -0.076 | 0.8818 | 0.03300638 | -0.074 | 0.8225 | 0.032137792 |
| RO_So_1 | RO_Somes | -0.221 | 0.8191 | 0.09597908 | -0.198 | 0.667 | 0.085990307 |
| RO_So_1 | RO_Somes | -0.152 | 0.7218 | 0.06601276 | -0.149 | 0.6525 | 0.064709878 |
| RO_So_2 | RO_Somes | -0.213 | 0.6687 | 0.09250473 | -0.175 | 0.4784 | 0.076001534 |
| RO_So_2 | RO_Somes | -0.166 | 0.7292 | 0.07209288 | -0.246 | 0.5183 | 0.106836443 |
